# Nuclear instance segmentation and tracking for preimplantation mouse embryos

**DOI:** 10.1101/2023.03.14.532646

**Authors:** Hayden Nunley, Binglun Shao, David Denberg, Prateek Grover, Jaspreet Singh, Maria Avdeeva, Bradley Joyce, Rebecca Kim-Yip, Abraham Kohrman, Abhishek Biswas, Aaron Watters, Zsombor Gal, Alison Kickuth, Madeleine Chalifoux, Stanislav Shvartsman, Lisa M. Brown, Eszter Posfai

## Abstract

For investigations into fate specification and cell rearrangements in time-lapse images of preimplantation embryos, automated 3D instance segmentation of nuclei and subsequent nuclear tracking are invaluable. Often, the images’ low signal-to-noise ratio and high voxel anisotropy and the nuclei’s dense packing and variable shapes limit the performance of many segmentation methods, while subsequent tracking of nuclear instances is complicated by low frame rates and sample movements. Supervised machine learning approaches can radically improve segmentation accuracy and enable easier tracking, but they often require large amounts of difficult-to-obtain annotated 3D data. Here we report a novel mouse line expressing near-infrared nuclear reporter H2B-miRFP720. H2B-miRFP720 is the longest wavelength nuclear reporter in mice and can be imaged simultaneously with other reporters with minimal overlap. We then generate a dataset, which we call BlastoSPIM, of 3D microscopy images of H2B-miRFP720-expressing embryos with ground truth for nuclear instance segmentation. Using BlastoSPIM, we benchmark the performance of seven convolutional neural networks and identify Stardist-3D as the most accurate instance segmentation method across preimplantation development. We then construct a complete pipeline for nuclear instance segmentation with our BlastoSPIM-trained Stardist-3D models and lineage tracking from the 8-cell stage to the end of preimplantation development (*>* 100 nuclei). Finally, we demonstrate BlastoSPIM’s usefulness as pre-train data for related problems, both for a different imaging modality and for different model systems. BlastoSPIM, its corresponding Stardist-3D models, and documentation of the full associated analysis pipeline are available at: blastospim.flatironinstitute.org.

## Introduction

During preimplantation development of the mouse embryo, two consecutive cell fate decisions set aside precursors of extraembryonic tissues from cells which will form the body of the embryo. Live images of embryos expressing fluorescently tagged proteins are particularly useful for learning the rules by which cells in the embryo dynamically interact with each other to specify these fates. However, deriving mechanistic insights from these images depends on extraction of quantitative information about cellular features, such as the position of each cell or the expression levels of specific proteins within each cell. Accurate segmentation of nuclei is a first step towards such a goal, as a cell’s nucleus is a good proxy for cell position relative to its neighbors and can contain information about cell-fate-specifying protein expression. To quantify these features, the segmentation must not only classify each voxel as foreground (i.e., belonging to nuclei) or background, but also assign each “instance” (i.e., nucleus) with a distinct label (S1 Fig).

Studying the dynamics of development requires instance segmentation not for a single frame, but for a (3+t)-D series of images of a developing embryo. To observe both fate decisions in preimplantation embryos, these movies start at the early morula stage (8-cell embryo) and end at the late blastocyst stage (*>*100-cell embryo), encompassing approximately 48 hours of development. Acquisition of a time lapse at sufficient spatial and temporal resolution to follow individual cells through 48 hours yields nearly 200 3D images (each composed of *≈*60 2D slices), containing a total of *≈*10,000 individual instances of nuclei; thus, manual segmentation of every instance in every frame is not feasible. Although classical image analysis methods have had success in automated nuclear segmentation [1–3], these methods often require high signal-to-noise ratio (SNR) images and tuning of parameters by hand. Shallow-learning methods, such as ilastik, offer an alternative solution for instance segmentation [4]; however, since these methods have relatively few trainable parameters, their performance saturates as the training set’s size grows [4]. Supervised deep-learning methods have many trainable parameters; thus, the performance of these networks benefit greatly from large ground-truth sets, which allow the networks to learn salient features. Relative to classical and shallow-learning methods, deep-learning methods often generalize better across biological conditions and microscopy types [5].

Deep-learning methods for 3D instance segmentation of nuclei differ considerably in terms of architecture, number of trainable parameters, and post-processing steps. Because of these differences (see S1 Table), it is difficult to know a priori which method will segment nuclei most accurately for any biological system of interest. To answer this question, ground-truth data is needed to a) train each network on relevant image annotations and b) to comprehensively test network performance by quantifying overlap between each instance in the ground-truth test set and each instance in the model output.

A study by Tokuoka *et al*. documented one of the first attempts to compare the performance of different deep-learning methods on a ground-truth dataset of nuclear instance segmentation in preimplantation mouse embryos [6]. Their ground-truth dataset, of nuclear centroids and semantic segmentation, spans from the 2-cell stage to at most the 53-cell stage (S2 Table) and enabled state-of-the-art performance for QCANet on early stages of development, up to approximately the 16-to-32-cell stage. The deterioration in performance of their model for later stages of development is in part due to the scarcity of training data past the 32-cell stage. Tokuoka *et al*.’s study demonstrated a clear need for nuclear instance segmentation that would perform accurately up to the end of preimplantation development in live images. For example, studying the first fate decision in mammalian preimplantation development – which differentiates those cells on the surface of the embryo (the trophectoderm, or TE) from those on the inside (the inner cell mass, or ICM) – requires accurate segmentation from the 8-cell stage to the *≈*64-cell stage [7]. Studying the next fate decision, in which ICM cells differentiate into epiblast and primitive endoderm cell populations that spatially segregate, requires accurate segmentation for even later stages (*>* 100 nuclei).

To this end, here we first generate a mouse line that expresses a near-infrared nuclear reporter H2B-miRFP720. H2B-miRFP720 is well suited for live imaging due to its reduced phototoxicity and its lack of spectral overlap with reporters in the visible range. Then, we generate a dataset, called BlastoSPIM (1.0), of light-sheet images of H2B-miRFP720-expressing preimplantation embryos with corresponding ground-truth for nuclear instance segmentation. This dataset is the largest of its kind. Historically, large ground-truth datasets have played a key role in enabling scientific progress. For example, by allowing researchers to focus on the methods rather than the data collection, annotation, and evaluation design, ImageNet [8] played an instrumental part in the rapid advances in object recognition. Our dataset for 3D nuclear instance segmentation similarly provides a foundation for evaluating both new and existing methods for nuclear instance segmentation. We use this dataset – that extends from the 2-cell stage to the *>*100-cell stage – to train and test seven different deep-learning methods, including Cellpose, Stardist-3D, RDCNet, U3D-BCD, UNETR-BCD, QCANet, and ELEPHANT. The Stardist-3D model, trained on BlastoSPIM 1.0, achieves state-of-the-art performance, detecting nuclei with high accuracy in early to mid-stage preimplantation embryos, even with low SNR images. Next, to improve segmentation accuracy at later embryonic stages, we generate a new ground truth dataset, termed BlastoSPIM 2.0, on blastocyst embryos and show that Stardist-3D trained on this dataset achieves similarly high nuclear segmentation accuracy even for embryos with *>*100 cells. By using the two (early and late) Stardist-3D models, we develop a complete pipeline for automatic segmentation, segmentation correction, nuclear centroid registration and lineage tracking. This pipeline, by enabling the tracking of nuclei from the 8-cell stage to the *>*100-cell stage, provides insight into the dynamics of nuclear volumes and nuclear shapes with respect to cell cycle and cell fate (ICM-TE). We close by demonstrating that our ground truth dataset and corresponding models aids nuclear segmentation in other model systems (intestinal organoids [9] and *Platynereis dumerilli* embryos [10]) as well as in data collected by other imaging modalities (spinning disk confocal).

## Materials and methods

### Transgenic mouse line generation

The H2B-miRFP720 transgenic mouse line was generated by targeting the TIGRE locus using the 2C-HR-CRISPR method [11]. Two targeting plasmids were constructed with InFusion cloning (Takara Bio), one consisting of 5’ and 3’ homology arms (each 1kb in length), surrounding H2B-miRFP720 driven by a CAG promoter and flanked by rabbit beta globin polyA sequence; the second construct contained an additional ORF-2A preceding H2B-miRFP720 flanked by a bGH polyA sequence. A single guide RNA (sgRNA) designed using CRISPOR [12] was used to target the TIGRE locus: CAUCCCAAAGUUAGGUGUUA (Synthego). CD1-IGS mice (Charles River strain 022) were used as embryo donors. Briefly, female CD1-IGS were superovulated at 5-7 weeks of age using 7.5IU PMSG (Biovendor) administered by IP injection followed by 7.5IU HCG (Sigma) by IP injection 47 hours post PMSG. Super ovulated females were mated to CD1-IGS stud males and checked for copulatory plug the following morning.

Cytoplasmic microinjection of 2-cell embryos was performed as previously described [11, 13]. Briefly, embryos were harvested at the 2-cell stage on E1.5 by flushing the oviducts with M2 Media (Cytospring) and each cell was microinjected with 100ng/ul Cas9 mRNA (made by IVT (mMESSGAE mMACHINE SP6 transcription kit, Thermo Fisher) using Addgene plasmid 122948), 30ng/ul donor plasmid and 50ng/ul sgRNA, using a Leica Dmi8 inverted epifluorescent microscope, an Eppendorf Femtojet and a Micro-ePore (WPI). Embryos were immediately transferred into the oviducts of pseudopregnant female CD1 mice. N0 pups were identified using over-the-arm PCR primers (Fwd:tcagcctacctcaccaactg, and Rev:ccccatcgctgcacaaaata) and outcrossed to CD1-IGS mice. N1 animals were genotyped using the same primers and the transgene was Sanger sequenced. The N1 generation was further outcrossed twice before incrossing the line to obtain homozygous mice. Homozygous and heterozygous offspring were distinguished using a wild-type PCR of the TIGRE locus (TIGRE WT Fwd:CTTTCCAGTGCTTCCCCAAC and TIGRE WT Rev: CCCTTTCCCAAGTCATCCCT).

The first mouse line showed decreasing levels of H2B-miRFP720 fluorescence during preimplantation development, while the second ORF-2A-H2B-miRFP720 mouse line showed ubiquitous high expression throughout. Therefore, the ORF sequence was deleted in 2-cell embryos isolated from this mouse line using the following sgRNAs: GGUGACGCGGCGCUGCUCCA and CAUGCCCAUUACGUCGGUAA, resulting in a truncated ORF with a functional 2A peptide. Founders and subsequent generations were established from this line, herein referred to as the H2B-miRFP720 mouse line, and ubiquitous H2B-miRFP720 fluorescence was confirmed once again in embryos. The sequence of the H2B-miRFP720 transgene can be found in S1 File. Other transgenic mouse strains used in this study include Cdx2-eGFP [14], mT/mG [15] and YAP-emiRFP670 [16].

### Dataset Acquisition

Embryos were obtained from naturally mated or superovulated H2B-miRFP720 females mated to either wild-type (CD1) or H2B-miRFP720 males. For the demonstration of four-color imaging *YAP-emiRFP670; Cdx2-eGFP* females were mated to *H2B-miRFP720; mT/mG* males. Embryos were isolated at E1.5 (2-cell), or E2.5 (8-cell) in M2 media and were cultured in Embryomax KSOM (Sigma-Aldrich) under paraffin oil (Life Global Paraffin Oil - LGPO from Cooper Surgical) in a V-shaped imaging chamber at 37°C, with 5% O_2_ and 5% CO_2_. Images were acquired on an InVi SPIM (Luxendo/Bruker). For each fluorescent reporter, the following excitation lasers and emission filters were used: eGFP 488 nm laser, 497-554 BP filter; tdTomato 561 nm laser, 577-612 BP filter; emiRFP670 642 nm laser, 659-690 BP filter; miRFP720 685 nm laser, 700 LP filter. To limit light exposure to the embryo, we acquired a full 3D image of each embryo at 15-minute intervals, with 2.0 μm z-axis resolution and 0.208 μm x- and y-axis resolution. Typically, the embryos were imaged from the 8-cell stage until the 64-cell stage or to the *>*100-cell stage, resulting in a duration of 48 hours or more. Raw time-lapse images were compressed to keller-lab-block (klb) format, on the fly. Blastocyst images used in Figure 6(A-C) were acquired using a W1 spinning disk confocal with SoRa on an inverted Nikon Ti2 with Hamamatsu ORCA-FusionBT, with a 20x (NA=0.75) air objective.

### Dataset Annotation

Raw 3D images of developing embryos were manually annotated using AnnotatorJ, an ImageJ plugin that supports both semantic and instance annotation. Images were loaded into the tool as Z stacks in .tiff format. For all images, brightness and contrast were adjusted by using the ‘auto’ and ‘reset’ functions in ImageJ. ‘Instance’ was selected as the annotation type. For each nucleus, the top or the bottom slice was found by comparing consecutive Z slices, and a contour was drawn for every slice that contained the nucleus. The coordinates of the regions of interest (ROIs) enclosed by the contours were then saved in an individual file. After each instance was annotated, the contours were overlaid on the image to distinguish the instance from unannotated ones. Five individuals annotated, and an expert checked for annotation errors, via a custom MATLAB code, before incorporation into the dataset.

Annotation of chromatin signal in mitotic cells, particularly those in metaphase and anaphase, presents unique challenges. Because our nuclear reporter H2B-miRFP720 is a tagged histone, chromatin condensation makes the fluorescent signal bright and often irregularly shaped. During metaphase, the ”nuclear” instance was annotated as contiguous, and its contour in each z-slice was drawn to closely match the shape of the signal. In anaphase and telophase, two instances (with distinct instance labels) were drawn and made to conform closely with the boundary of the bright condensed signal. We carefully annotated the H2B signal in metaphase, anaphase and telophase because it is important for the trained network to segment these nuclei well: the shape – particularly the orientation of the metaphase plate – and volume can be particularly informative for the assignment of daughter nuclei to mother nuclei in time lapse images (see Description of Semi-automated Nuclear Tracking Methods).

### Dataset Characteristics

The BlastoSPIM 1.0 dataset includes 573 fully annotated 3D images of nuclei in mouse embryos, each manually curated for annotation. Across all images, there are 11708 individual nuclear instances annotated and 116 annotated polar bodies. Not all of these 3D images come from different time series. For example, for one developing mouse embryo, we annotated 89 consecutive time-points, and for another embryo, we annotated 100 consecutive time-points. The total number of distinct embryos imaged and annotated is 31.

Aside from diversity in developmental stage, the embryos in this dataset express different H2B-miRFP720 alleles (see details in mouse line generation) and were also imaged with different laser intensities. This diversity in SNR allows us to test whether model performance degrades significantly as SNR decreases. We quantify SNR in our case by calculating mean foreground intensities and mean background intensities. We report the distribution of SNRs, one point for each fully annotated 3D image, as the difference between mean foreground and mean background in (S2 Fig). For comparison, the background intensity – in gray values – typically has a mean of 118 and a variance of 10-14.

The BlastoSPIM 2.0 dataset consists of 80 annotated images of late-stage embryos. This set includes 6628 nuclear instances. Late blastocysts from this dataset with the lowest SNR values (S2 Fig) were selected and incorporated into the existing low SNR set. The final number of annotated images in BlastoSPIM 1 + 2 is 653, and the number of annotated nuclear instances is 18336.

In addition, to demonstrate that the models trained on BlastoSPIM perform well also for images acquired via other modalities, we annotated the nuclei in 10 different embryos imaged on a spinning disk confocal microscope (Figure 6(A-C)). These ten images range from the 30-cell stage to the 61-cell stage and contain 461 nuclei in total.

### Dataset Splits and Evaluation Metric

When splitting our dataset into a training set, a test set, and a validation set, our main objective was to quantify how model performance varies as a function of both developmental stage and SNR. For BlastoSPIM 1.0 we created two separate test sets, one for low SNR and one for moderate SNR, each of which contained a diversity of developmental stages. We define “low SNR” and “moderate SNR” by comparing the mean foreground intensity to the mean background intensity. The “low SNR” images all have a mean foreground intensity which is at most 134 gray values, approximately 15 gray values above the typical mean background intensity. For reference, the background intensity – in gray values – typically has a variance of 10-14 (S2 Fig). Within both the moderate SNR and the low SNR sets, we group annotated embryos based on their developmental stage, estimated by the number of nuclei (*i.e*., *≈*8-cell, *≈*16-cell, *≈*32-cell, *≈*64-cell, *>*100-cell). Each set deliberately contains more images from earlier stages than later stages so that the total of number of nuclei per developmental stage is at least partially equalized across stages. From the BlastoSPIM 2.0 dataset, 8 embryos from various stages were used as a test set, as early as the 48-cell stage and as late as the 107-cell stage. The remainder of the data, 72 annotated embryos, were either for validation or training. The exact breakdown is specified at blastospim.flatironinstitute.org.

To evaluate how well the models performed on the test sets, we computed the intersection-over-union (IoU) between the models’ segmentation and the ground truth. We considered an instance in the models’ segmentation to match an instance in the ground truth if the IoU between the two was at least 0.5. We also provide how performance varies as a function of this IoU threshold (S3 Fig, S4 Fig). We also calculate the IoU between its ground-truth instance and its matched model-inferred inference; each unmatched ground-truth instance contributes an IoU of zero to the average. As above, the IoU must be at least 0.5 for each match. To compute how well the predictions fit the ground truth, we compute the panoptic quality [17]: 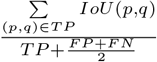, where (*p, q*) represents a match between predicted object *p* and ground-truth object *q. TP, FP*, and *FN* denote true positives, false positives, and false negatives, respectively.

The train-test-validation split for all the models is specified on the BlastoSPIM website.

### Segmentation Correction Methods

To achieve accurate lineage tracking, even infrequent segmentation errors need to be corrected. These errors include oversegmentation, undersegmentation, or misclassification of background noise as cells. To manually rectify these errors, we employed an enhanced version of the ImageJ plugin, AnnotatorJ [18].

AnnotatorJ provided users with an intuitive interface to inspect and identify segmentation errors by overlaying the segmented ROIs onto the original image. To cater to the specific needs of 3D image analysis, we introduced various enhancements to AnnotatorJ (v1.6). Briefly, AnnotatorJ v1.6 allowed for the display of ROIs in each frame of a 3D image, enabling users to navigate seamlessly through frames and address errors in each frame individually. This functionality was made possible by incorporating operations to add or remove regions, modify boundary positions, and correct misclassified areas manually. A ”generate mask” operation was added to easily generate the labeled mask image once corrections were completed. We implemented color coding of ROIs for easier identification and generation of a final corrected labeled mask image. Additionally, we introduced support for loading compressed KLB files, for efficient data handling.

### Registration Methods

Under normal imaging conditions, the embryo can rotate and translate within the microscope field of view. While global positions of cells in consecutive frames may change, their relative positions are unlikely to change significantly. To achieve positional consistency over time, we use the coherent point drift (CPD) algorithm to register pairs of frames in sequence [19]. CPD is a point set registration algorithm which takes a probabilistic approach to aligning two sets of points. Each point set is represented by a gaussian mixture model (GMM), and a transformation function is learned which maps the centroids of the first point set onto the centroids of the second set. For each pair of frames, we take the corresponding point sets to be the centroids of the distinct instances in the instance segmentation. We also restrict the transformation function to be a rotation and translation.

We found that CPD tended to converge to local minima on our data depending on the initial rotation guess. Because the point sets were small and thus the CPD execution time was low, the correct registration could be found quickly by running many CPD trials with randomly chosen initial rotations. The final registration was chosen as the trial which minimized the GMM covariance parameter. This covariance parameter acts as a length-scale which tends to be large for misaligned point sets and small for aligned point sets.

### Nuclear Tracking

After the error correction and registration steps, we perform semi-automated lineage tracking on the registered nuclear segments. Our tracking algorithm constructs the lineage tree sequentially, at every iteration matching nuclei to their predecessors in the previous time frame. We build our algorithm on the previously published effort [20] that was based on nearest neighbor association between nuclear centroids at adjacent time frames and, in the case of mitosis, searching for pairs of daughters with minimal distance to the mother. However, we found that while this method can be successful in tracking nuclei during interphase, the slower frame rate of our samples is an obstacle to the success of the algorithm in correctly identifying division events. Therefore, we introduced additional steps comparing nuclear volumes for every matched pair of nuclei to prune incorrect tree edges and identify mother-daughter triples, with the option for manual correction. This approach takes advantage of the observed differences in nuclear volume between daughter nuclei and their mother due to splitting of condensed chromatin. More precisely, for every pair of frames, we perform several steps. In the steps below, regardless of whether a division event occurred between times *t* and *t* + 1, we refer to nuclei at time *t* as mothers (or parents, interchangeably) and nuclei at time *t* + 1 as daughters. Only if a mother at time *t* has two daughter nuclei at *t* + 1 has a division event occurred in the tree.

#### Step 1

We start by matching every label at time *t* + 1 with its nearest neighbor at time *t* (using euclidean distances between centroids), thus identifying prospective parents for every label. The matches can be viewed as the edges that are added to the tree at height *t*. For these initial edges, we compare the volumes of the two matched nuclei and first retain the edges that represent one-to-one mappings without *significant volume disbalance* (defined by the daughter-mother volume ratio larger than a user-defined threshold; heuristically, we set the default value to be 2/3). The rest of the edges represent many-to-one daughter-mother mappings. For every prospective mother, we identify how many of the prospective daughter nuclei are large enough, i.e., do not demonstrate significant volume disbalance to the mother (defined above). If there is just one large daughter identified, we retain this connection pruning all the rest; otherwise, we prune all the edges for this prospective mother. Thus, all the retained matches potentially represent the same nucleus which slightly changed its position between times *t* and *t* + 1.

#### Step 2

At this point, the nuclei at times *t* and *t* + 1 that belong to the edges retained at Step 1 are removed from consideration. The remaining nuclei at time *t* + 1 undergo the next round of nearest neighbor mapping to time *t*. At this stage, one-to-one mappings are retained. Now we aim to identify for mitotic triples. To do this, we search for prospective parents (time *t*) that were mapped to exactly two nuclei at time *t* + 1. We retain both edges for such a triple if there is significant daughter-mother volume disbalance (see definition in Step 1) for both prospective daughters. The mitotic triple criterion includes an option to check for the centroid of the daughters to be close enough to the prospective parent.

#### Step 3

All the remaining connections from Step 2 represent the many-to-one mappings that do not satisfy the criterion for the mitotic triple. We attempt to resolve such conflicting matches by using second nearest neighbors. If this procedure does not identify a plausible mitotic triple based on the criterion described in Step 1, we manually identify the correct mitotic triple and edit the lineage tree by using the rmedge and addedge functions in MATLAB.

### Construction of the Lineage Trees from the 8-cell stage to the *≈* 100-cell stage

The initial segmentation was produced using the “early embryo” Stardist-3D model for frames 1 through 145, at which point the embryo’s cell count reached 64. For the remainder of the movie, ending at frame 210, we switched to the “late blastocyst” Stardist-3D model as it achieves a higher accuracy in later stages and reduces the number of error corrections needed. Following segmentation, we used our enhanced AnnotatorJ tool to hand-correct errors and began tracking using our semi-automatic lineage construction algorithm. The process of lineage construction provides temporal context which often reveals errors in the segmentation which may be non-obvious when viewing a segmented frame in isolation. For this reason, performing hand-correction and lineage construction in parallel is useful. After building the lineage tree, inner cell mass (ICM) and trophectoderm (TE) fates were assigned based on nuclear shape (TE nuclei are typically more flat) and position (TE nuclei are closer to the surface, while ICM nuclei are positioned deeper in the embryo).

From the beginning of the 32-cell stage to the 64-cell stage, the estimated volume of each nucleus was computed by averaging the predictions of the early and late models. More specifically, since only one of these two models was corrected per time point for the purpose of the lineage tree, we computed matches between the corrected segmentation (from either the early or late model) and an uncorrected segmentation from the other model. If a match of sufficiently high IoU was identified, then the volumes of the nuclear instance in the two models were averaged. If no sufficiently matching instance was found, then only the volume of the instance in the corrected segmentation was used. S5 Fig demonstrates that the model-predicted instances have volumes which agree well with those of the ground-truth matches; thus, for Figure 5 we used model-predicted nuclear volumes as a reasonable proxy for the ground-truth volumes (which can only be obtained by full manual annotation – not done for these lineage trees).

### Quantifying nuclear volume and aspect ratio

For the quantification of nuclear aspect ratio, we use the procedure – based on calculating the moment of inertia tensor – described in [21]. The quantification of volume was obtained by multiplying the number of vocels in the instance with voxel volume. For Figure 5(D,E), to reduce noise in estimated nuclear volume, each maximal volume (*V*_*max*_) per cell stage (8-to-16, 16-to-32 or 32-to-64) was calculated by averaging the nuclear volume for 90 minutes preceding the nuclear volume peak (see Figure 5(B)).

For calculating maximal nuclear volumes in Figure 5(D,E), the 8-to-16 and 16-to-32 divisions are unambiguously defined because they occur in two temporally distinct rounds. On the other hand, because cell cycle asynchrony increases with developmental time, the definition of a 32-to-64 cell division is potentially ambiguous: namely, some 16-to-32 divisions have not happened even as some daughters of other 16-to-32 divisions have themselves divided (see Figure 5(A)). For each tree, along a path from the root to the leaves, we define the 32-to-64 cell divisions as those which – without any intervening divisions along the lineage path – follow a 16-to-32 division. For the purposes of Figure 5(D,E), we only plot the 32-to-64 divisions which result in two progeny which both survive until the end of tree and divide at least once more. We make this choice because those are the only cases in which the quantities *V*_*D*1,*max*_ + *V*_*D*2,*max*_ and *V*_*D*1,*max*_ *− V*_*D*2,*max*_ are well-defined.

## Results

### 0.1 Establishment of a near-infrared nuclear reporter mouse line

Multicolor imaging is key to simultaneous recording of morphogenesis and cell fate specification. To enable visualization of cell nuclei in concert with various other molecular markers, which are typically tagged with green, red or far-red fluorescent proteins, we generated a novel spectrally distinct near-infrared nuclear mouse line expressing H2B-miRFP720 (Fig 1(A)-(B)). First, using 2C-HR-CRISPR [11] we targeted CAG H2B-miRFP720 to the TIGRE locus [12]. Early preimplantation embryos from this line showed uniform H2B-miRFP720 expression; however, by the mid blastocyst stage significant dimming of the fluorescent signal was noted, even in freshly isolated embryos (data not shown). A second mouse line harboring CAG ORF-2A-H2B-miRFP720 in the TIGRE locus however did not exhibit the same dimming issue, and rather showed a slight increase in H2B-miRFP720 intensity during preimplantation development. We therefore used two sgRNAs to delete the ORF-2A with Cas9 in this line, resulting in a CAG H2B-miRFP720 line (hereafter referred to as the H2B-miRFP720 mouse line) with bright reporter expression across all preimplantation stages (Fig 1(A)-(B)). miRFP720 can be readily multiplexed with far-red fluorescent reporters such as emiRFP670 or the Halo-tag visualised with the JF646 dye, therefore this mouse line allows simultaneous imaging of up to four different reporters in mouse embryos (Fig 1(C)). Furthermore, the long wavelength used for its detection makes H2B-miRFP720 ideal for deep-tissue imaging and results in reduced phototoxicity.

**Fig 1.**
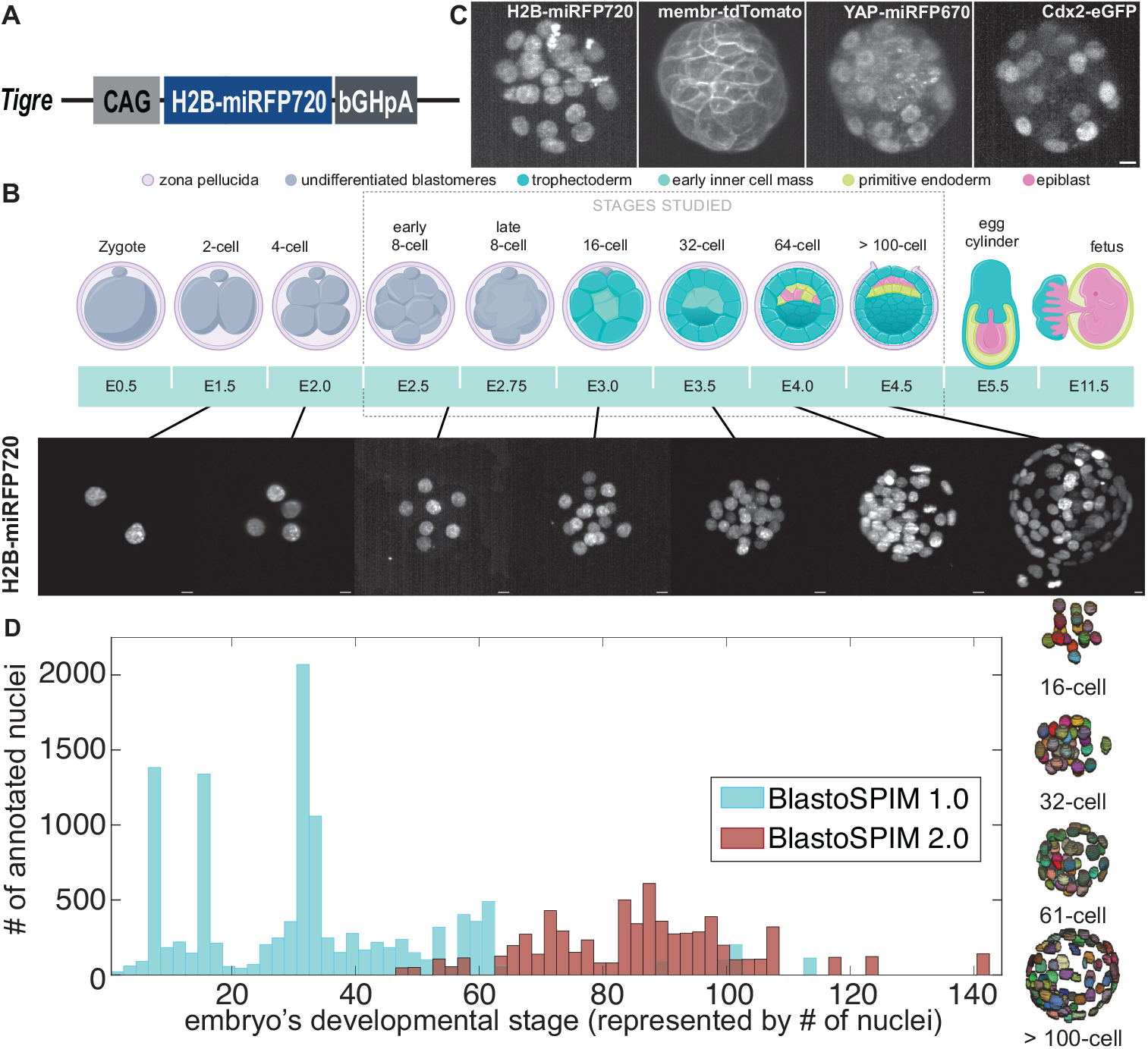
BlastoSPIM datasets, ground truth of nuclear instance segmentation for embryos expressing a novel near-infrared nuclear marker. (A) Schematic of targeted TIGRE locus with the CAG-H2B-miRFP720 insert. (B) Top: cartoon of preimplantation development in the mouse. After fertilization, the zygote undergoes cleavage divisions. At the 8-cell stage, compaction and polarization occur. By the 32-cell stage, a subset of cells called the trophectoderm (TE) form the embryo’s surface; the remaining cells form the inner cell mass (ICM). The ICM cells begins to pattern into two fates, primitive endoderm (PE) and epiblast (EPI), by the 64-cell stage; by implantation, around the *>* 100-cell stage, the two inner fates are spatially segregated. Bottom: Maximum-intensity projected images – acquired with SPIM – of preimplantation embryos expressing H2B-miRFP720 at different developmental stages. Scale bar: 10 *μm*. (C) Preimplantation embryo expressing four spectrally distinct fluorescent reporters: H2B-miRFP720, mTmG, YAP-emiRFP670, Cdx2-eGFP. Maximum intensity projections of images acquired with SPIM. Scale bar: 10 *μm*. (D) Histogram of number of nuclei per embryonic stage (represented by embryo cell number) for both BlastoSPIM 1.0 (blue, used for initial benchmarking of methods) and BlastoSPIM 2.0 (red, used for extending accurate segmentation to later stages). For four embryos from different stages, the ground truth of nuclear segmentation are shown.

### 0.2 A novel ground-truth dataset of preimplantation mouse embryos for comparing nuclear-segmentation methods

Using selective plane illumination microscopy (SPIM) we acquired 3D live images of H2B-miRFP720-expressing preimplantation embryos at various developmental stages. We created a new ground-truth dataset with full 3D nuclear instance segmentation. This dataset, which we call BlastoSPIM 1.0 (concatenation of blastocyst and SPIM), is one of the largest and most complete of its kind (S2 Table) with more than 570 high-resolution, light-sheet images with approximately 12,000 nuclei are annotated, spanning all preimplantation stages (Fig 1(D)) (for details, see Dataset Characteristics and S2 Fig). The quality, detail, and size of the BlastoSPIM dataset makes it unique relative to other publicly available ground truth datasets for nuclear instance segmentation (S2 Table).

To quantitatively illustrate the challenges posed by densely packed nuclei for instance segmentation, using BlastoSPIM 1.0, we calculated how nucleus-to-nucleus distances change from the 16-cell stage to the *>*100-cell stage. The surface-to-surface distance between nearest-neighbor nuclei has a median of 6.0 *μm* at the 16-cell stage, 2.9 *μm* at the 32-cell stage, 1.8 *μm* at the 64-cell stage, and *≈* 0.5 *μ* at the *>*100-cell stage (S6 Fig). This decrease in nearest-neighbor distance, with an increasing number of nuclei having a *<* 1*μm* nearest-neighbor distance with successive developmental stages, is not accompanied by a comparable decrease in nuclear size (S6 Fig); thus, instance segmentation is expected to be considerably more challenging as development progresses.

Additionally, the challenge for instance segmentation is due not only to nucleus-to-nucleus juxtaposition, but also to characteristics of image acquisition. For example, live images often have low SNRs because the exposure of embryos to light has to be limited to prevent phototoxicity [22]. Moreover, the sample is imaged along a single axis (z-axis, by convention), resulting in voxel anisotropy – poorer z-resolution than xy-resolution. Our ground-truth dataset contains a range of SNR values (S2 Fig) and has a voxel anisotropy of approximately 10. In summary, because of its size as well as its diversity in developmental stage and SNR, our dataset of manually annotated 3D instances of nuclei is uniquely suited to interrogate the performance of any segmentation method – for achieving accurate nuclear instance segmentation in preimplantation mouse embryos.

### 0.3 Benchmarking of seven instance segmentation methods on BlastoSPIM 1.0 reveals superior performance of Stardist-3D

We used our ground-truth dataset to compare seven instance segmentation networks (in S1 Table), including Cellpose [23], Stardist-3D [24], RDCNet [25], U3D-BCD [26], UNETR-BCD [27], ELEPHANT [28], and QCANet [6]. These methods span a variety of network architectures, from those including recurrent blocks or transformers to more conventional U-Nets. They also represent the instances in different ways. For example, Stardist-3D computes a set of distances to the boundary, while Cellpose predicts gradients that are tracked to the instance center.

We trained each model with data from 482 3D images of embryos from BlastoSPIM 1.0 and then evaluated on a test set composed of moderate SNR data. To interrogate stage-specific performance, we divided this test set into developmental stages such that it contained approximately 120 nuclei from each stage (*e.g*., more images from earlier stages than later stages). To benchmark each method, we compared the ground-truth instances and model-inferred instances by computing matches based on the intersection-over-union (IoU). A model-inferred instance is considered as a match to a ground-truth instance if the IoU is at least 0.5 (see different IoU cutoffs in S3 Fig). Based on this matching, we computed the *F*_1_ score, defined as 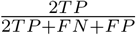, where TP, FP, and FN are the number of true positives, false positives, and false negatives, respectively (Figure 2(A)). We also quantified the accuracy of the model predictions based on the panoptic quality (see Dataset Splits and Evaluation Metric, (Figure 2(B)). As opposed to the *F*_1_ score, which simply counts matches in a binary way based on a threshold in IoU, the panoptic quality also depends on the sum of the IoUs that are above the specified IoU threshold.

**Fig 2.**
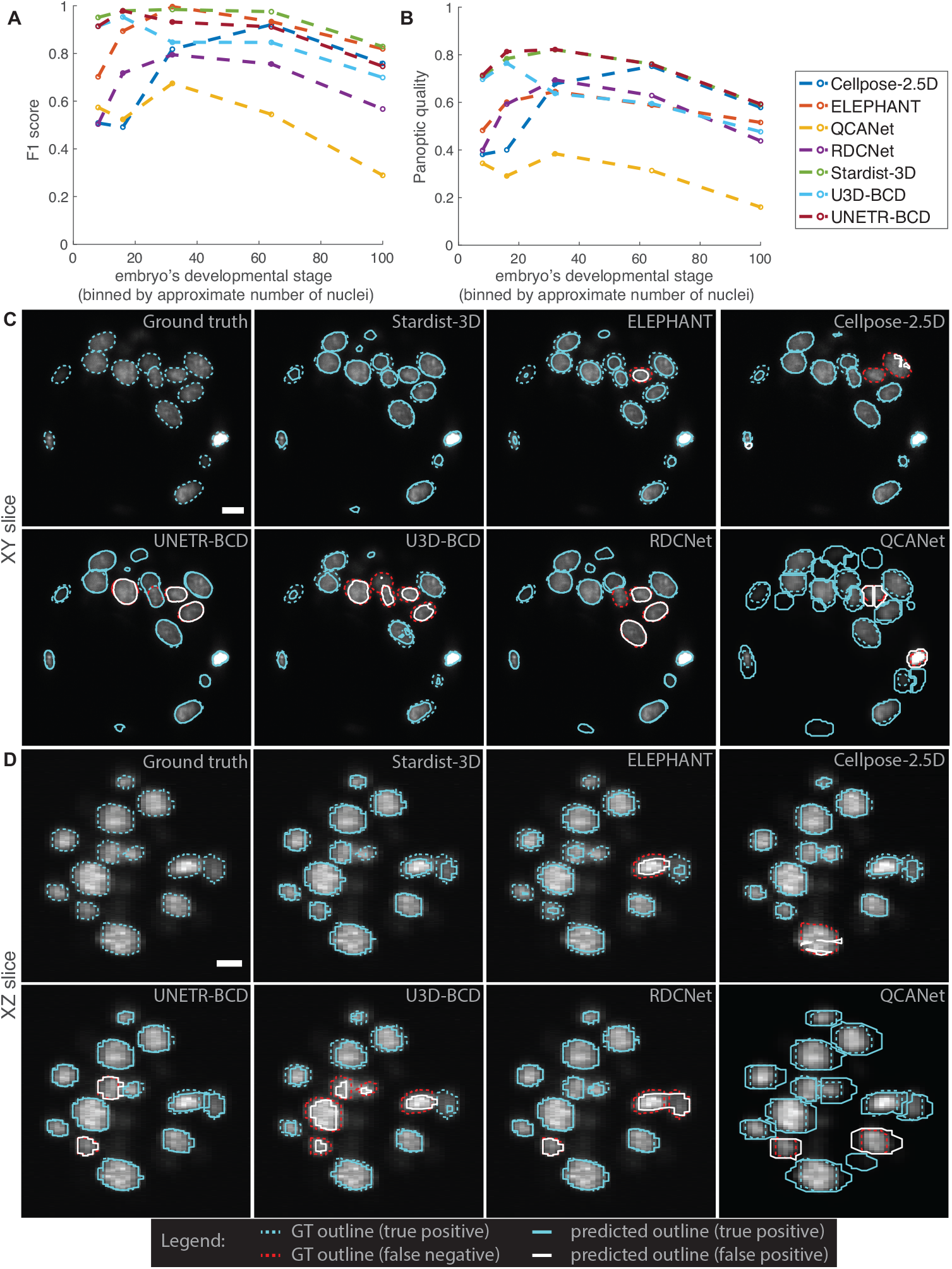
Evaluation of seven instance-segmentation networks trained on BlastoSPIM 1.0 across preimplantation developmental stages. (A,B) F1-score and panoptic quality, respectively, across developmental stage. See S3 Fig for evaluation at different IoU cutoffs. (C,D) Qualitative evaluation on a 60-cell embryo. Instance contours overlaid on a representative slice of the intensity image in xy and in xz, respectively. Each panel is labelled as either Ground-truth or according to the method evaluated. GT, predicted true positive, false negative and predicted false negative outlines are shown for each model (see legend in figure). False positives and false negatives are defined by comparing the 3D instance segmentation results rather than the results shown in a single 2D slice. Note that if the model predicted instance does not overlap sufficiently well with the ground truth instance, the result is a false positive paired with a false negative. In (C), the extra cyan outlines for QCANet are predicted instances which match with instances in nearby z-slices but over-extend in z. Scale bars: 10 *μm*. See S3 Table, S4 Table for evaluation of model failures for difficult cases in other test images.

Based on the *F*_1_ score, the Stardist-3D model outperformed all other methods across developmental stages (Figure 2(A)). From the 8-cell stage up to the 64-cell stage, the *F*_1_ score remained above 95 %. This is significantly higher than the state-of-the-art results on similar (confocal) data from preimplantation mouse embryos, particularly in embryos with *>* 32 nuclei [6]. By comparison to Stardist-3D’s strong *F*_1_ score across stages, the *F*_1_ score of the other methods depended more strongly on developmental stage (Figure 2(A)). For example, both UNETR-BCD and the related U3D-BCD method performed reasonably well at the 8- and 16-cell stages but were unable to detect several nuclei in later stages as nuclei became more densely packed. By contrast, the performance of Cellpose and RDCNet slightly improved from the 8-cell stage to the 64-cell stage, then decreased at the later developmental stages.

Stardist-3D also achieved the highest panoptic quality, approximately equal to that of UNETR-BCD, across developmental stages (Figure 2(B)). Despite UNETR-BCD’s low *F*_1_ as compared to both Stardist-3D and ELEPHANT, its high panoptic quality can be explained as follows: the IoUs for UNETR-BCD’s successful matches to the ground-truth instances are high even though it has fewer matches. By contrast, although ELEPHANT had a reasonably high *F*_1_ score for the 16-cell and later stages, the panoptic quality remained relatively low across stages. This is likely explained by ELEPHANT’s constraint that nuclei be represented only as ellipsoids; by contrast, Stardist-3D’s representation of nuclei as star-convex polyhedra approximates well the nuclear shapes in our ground-truth dataset S7 Fig.

In terms of both performance metrics, one method, QCANet, performed worse than the other methods. Two key factors likely contributed to this poor performance. First, QCANet makes the images isotropic by decreasing the xy-resolution and interpolating in z, and this coarser resolution likely complicates the prediction of the boundary between closely juxtaposed nuclei. Second, since QCANet uses centroid-based watershed, errors in the predicted centroid locations impact the predicted boundaries between instances and can give rise to the prediction of instances with unrealistic shapes.

Figure 2(C,D) shows qualitative results of these seven networks on an embryo with 60 nuclei based on two 2D image slices (one in xy, the other in xz). On this test image, Stardist-3D achieved the best F1-score by producing only one false negative (not shown in slice) and no false positives. Although ELEPHANT produced one instance per ground-truth instance, two of those instances overlapped poorly with the corresponding ground-truth; these resulted in two false negatives and two false positives. UNETR-BCD and Cellpose each produced approximately ten errors, including false positives and false negatives. The remaining three methods produced significantly more errors than the others on this test case. Below we summarize the typical errors made by each network.

U3D-BCD, UNETR-BCD, and RDCNet missed several nuclei due to under-segmentation – the merging of more than one nucleus into the same instance label (see Figure 2(C) for UNETR-BCD and Figure 2(D) for RDCNet). On the other hand, Cellpose often oversegmented nuclei into small spurious instances or missed a ground truth instance entirely (predicting no instance which overlaps; see Figure 2(C)). QCANet often predicted the right number of nuclear centroids, and thus avoided over- or under-segmentation, but if two predicted centroids fell within the same contiguous region of the semantic segmentation, the watershed-based post-processing often split the mask improperly (see two neighboring false positives in Figure 2(C)).

Some methods produced instances with shapes that are not consistent with the range of ground-truth nuclear shapes. For example, QCANet often overpredicted the nuclear sizes (see instances which extend across more z-slices than the ground-truth instances in Figure 2(D)). By contrast, Cellpose and RDCNet tended to produce some small false positives with highly irregular shapes (see small false positives in Cellpose panels in Figure 2(C-D)). U3D-BCD tended to produce instances with holes or gaps (see small holes in U3D-BCD instance in Figure 2(C)). Although models (like U3D-BCD or RDCNet) that make few hard assumptions about nuclear shape sometimes performed best on particularly difficult cases like mitotic nuclei or polar bodies (S3 Table, S4 Table), our results suggest that the methods which make biologically plausible assumptions about nuclear shape (ELEPHANT and Stardist-3D) often achieve higher *F*_1_ scores. In summary, our comprehensive benchmarking of different models provided insight into the strengths and weaknesses of each network and identified Stardist-3D as the best performing network for 3D images of live preimplantation embryos. Moreover, our large ground truth dataset can be used to test whether future neural network architectures can outperform Stardist-3D in accurately identifying the positions and shapes of nuclei in embryos.

### 0.4 Extending Stardist-3D’s segmentation accuracy up to the *>* 100-cell stage

Although Stardist-3D performed well up to the *≈* 64-cell stage (Figure 2(C-D)), its performance deteriorated at later stages. Therefore, we set out to improve accuracy by specifically training the network on late stage ground-truth data. We hand-annotated an additional 80 3D images of late stage embryos expressing H2B-miRFP720, containing more than 6600 nuclear instances – a data set we termed BlastoSPIM 2.0 (Fig 1(D)). We trained and validated a new Stardist-3D model based on 72 images of late blastocysts from BlastoSPIM 2.0 (termed ”late blastocyst model”).

The late blastocyst model outperformed the previous Stardist-3D model (hereafter referred to as the ”early embryo model”) from Figure 2 on test images of late blastocysts, and underperformed it on test images of embryos with fewer than 64 nuclei (Figure 3(A,B), S4 Fig, S8 Fig). In particular, the late blastocyst model performed better than the early embryo model in cases with closely juxtaposed nuclei (as seen in the ICM) (Figure 3(C), S9 Fig), while it often over-segmented polar bodies, which typically only appear in images of early embryos (S8 Fig). For comparison to these two models, we also trained a model on both BlastoSPIM 1.0 and 2.0 (”all stages model”) and found that this model’s performance fell between the performance of the other two Stardist-3D models (Figure 3(A,B), S4 Fig). As a result, at any developmental stage, one would achieve higher performance with the stage-specific models. We therefore transitioned from using the early embryo model to using the late blastocyst model when the embryo has *≈* 48 nuclei (in the transition between the 32-cell stage and the 64-cell stage).

**Fig 3.**
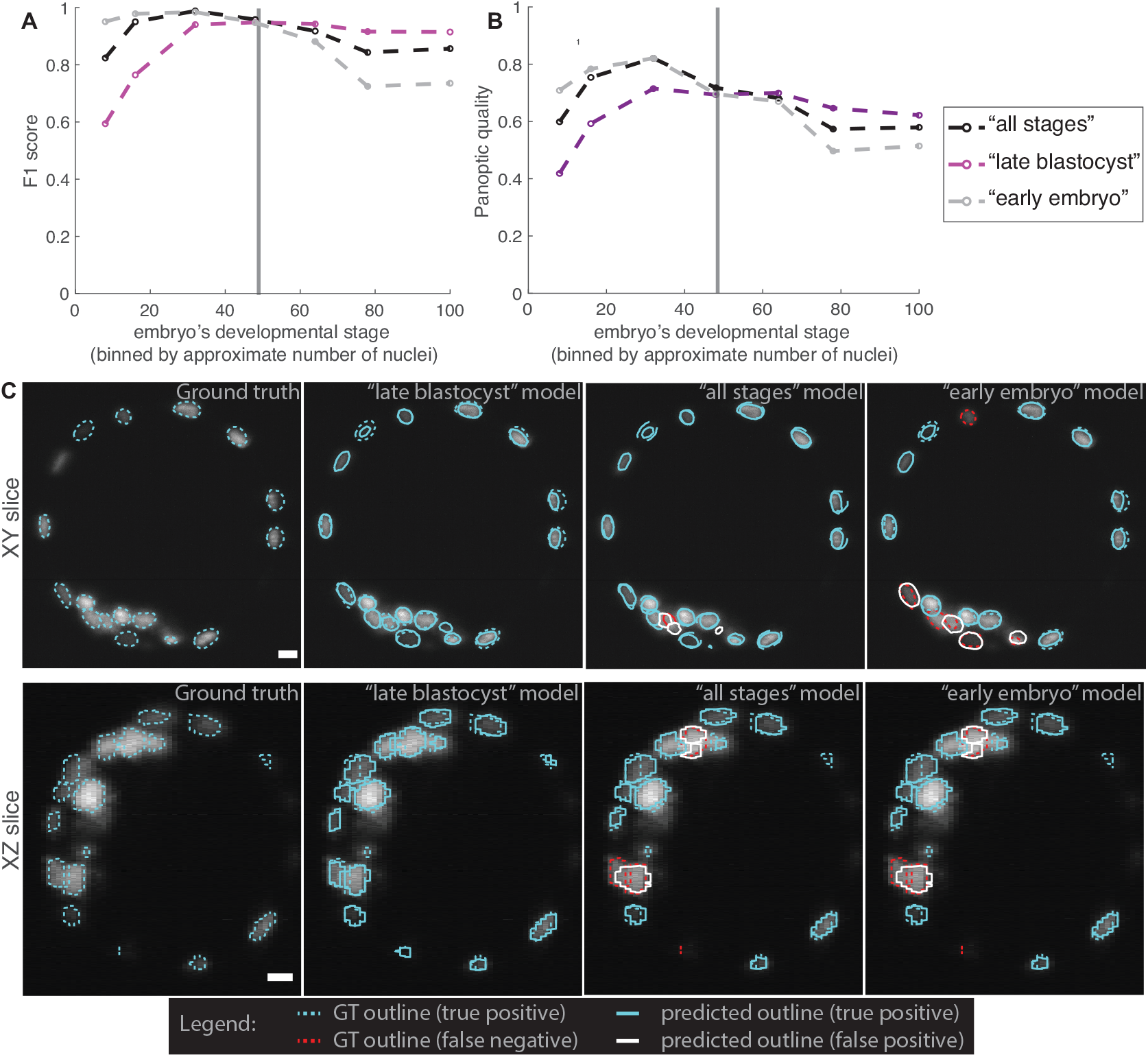
Comparing models trained on BlastoSPIM 1.0 (early embryo model), BlastoSPIM 2.0 (late blastocyst model), and on both 1.0 and 2.0 (all stages model). (A,B) F1-score and panoptic quality, respectively, for the moderate-to-high SNR test set. See S4 Fig for evaluation at different IoU cutoffs. Vertical line: approximate developmental stage at which to transition from the early embryo model to the late blastocyst model. (C) Qualitative evaluation on a 106-cell embryo. Instance contours overlaid on a representative slice of the intensity image in xy (top) and in xz (bottom), respectively. Each panel is labelled as either Ground-truth or according to the model evaluated. Scale bars: 10 *μm*. Note that if the model predicted instance does not overlap sufficiently well with the ground truth instance, the result is a false positive paired with a false negative. Relatedly, see S8 Fig for errors produced by inference of the late blastocyst model on early embryo data.

We next asked whether the early embryo model and late blastocyst model would perform well even for cases of very low SNR. We, thus, evaluated these Stardist-3D models on a more difficult test set – separate from the test set in Figure 2 – comprised of images with a very low SNR ratio (S9 Fig). Although only a third of the images in the training set for the early embryo model met our definition of low SNR (S2 Fig), the *F*_1_ score of the early embryo model for the low SNR test set was ⪆ 90 % for all stages up to the 64-cell stage. For later stages, the late blastocyst model outperformed the early embryo model by achieving an *F*_1_ score of *≈* 86 %. Since the images in our low SNR test represent some of the most challenging cases, where even human experts have difficulties in annotating instances (S9 Fig(D)), we expect our models to generalize well to long-term live-imaging datasets with low-to-moderate SNR.

### 0.5 BlastoSPIM-trained Stardist-3D models enable lineage tracking

Analyzing developmental dynamics requires tracking of individual cell lineages over time. We therefore developed a complete pipeline image analysis pipeline that integrated nuclear segmentation results from the BlastoSPIM-trained Stardist-3D models with lineage tracking. First, using light-sheet microscopy, we acquired a time series of 3D images of an H2B-miRFP720-expressing embryo, with one Z-stack acquired every 15 minutes. We automatically segmented the images by either the early embryo model or late blastocyst model, switching models around the 48-cell stage (Figure 4(A)). As accurate lineage tracking is contingent upon accurate segmentation, we hand-corrected remaining errors using AnnotatorJ [18]. AnnotatorJ overlays each segmented region of interest (ROI) onto the original image and provides an intuitive interface to edit, delete, or create ROIs. Since AnnotatorJ was originally designed for 2D images, we introduced several enhancements for corrections of 3D segmentations. Segmentation errors were corrected by removing false positives, adding missing instances, and editing over-segmented regions.

**Fig 4.**
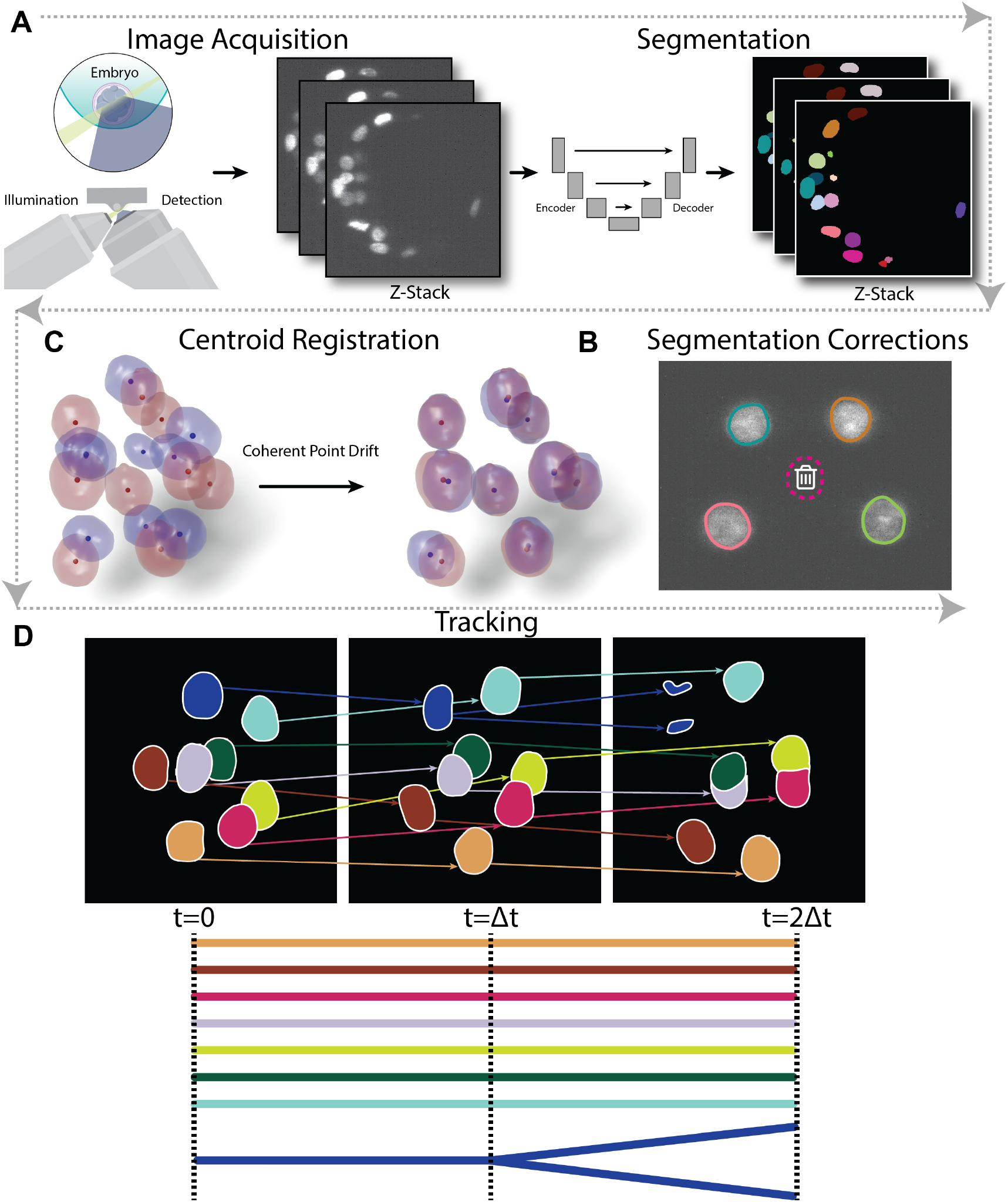
Analysis pipeline, from image acquisition to lineage tree construction. Gray arrows indicate order of steps in the pipeline. (A) Image acquisition and segmentation. Time series of 3D light-sheet images were segmented automatically using the early embryo and the late blastocyst Stardist-3D models. Green: light sheet used for illumination. Blue: emitted light is collected by the detection objective. (B) Segmentation corrections. Errors in the segmentation were hand-corrected by overlaying the segmentation with the raw image in each frame and removing false positives and/or adding instances which were missing. (C) Registration of nuclear segmentations. Using the corrected segmentation, consecutive pairs of frames are aligned spatially to account for rigid motion the embryo experiences during image acquisition. We employ the coherent point drift registration algorithm, operating on the instances’ centroids extracted from the segmentation. (D) Lineage tracking. After registration, a set of lineage trees was constructed (one for each of the nuclei in the first frame) by matching nuclear identities between pairs of consecutive frames.

**Fig 5.**
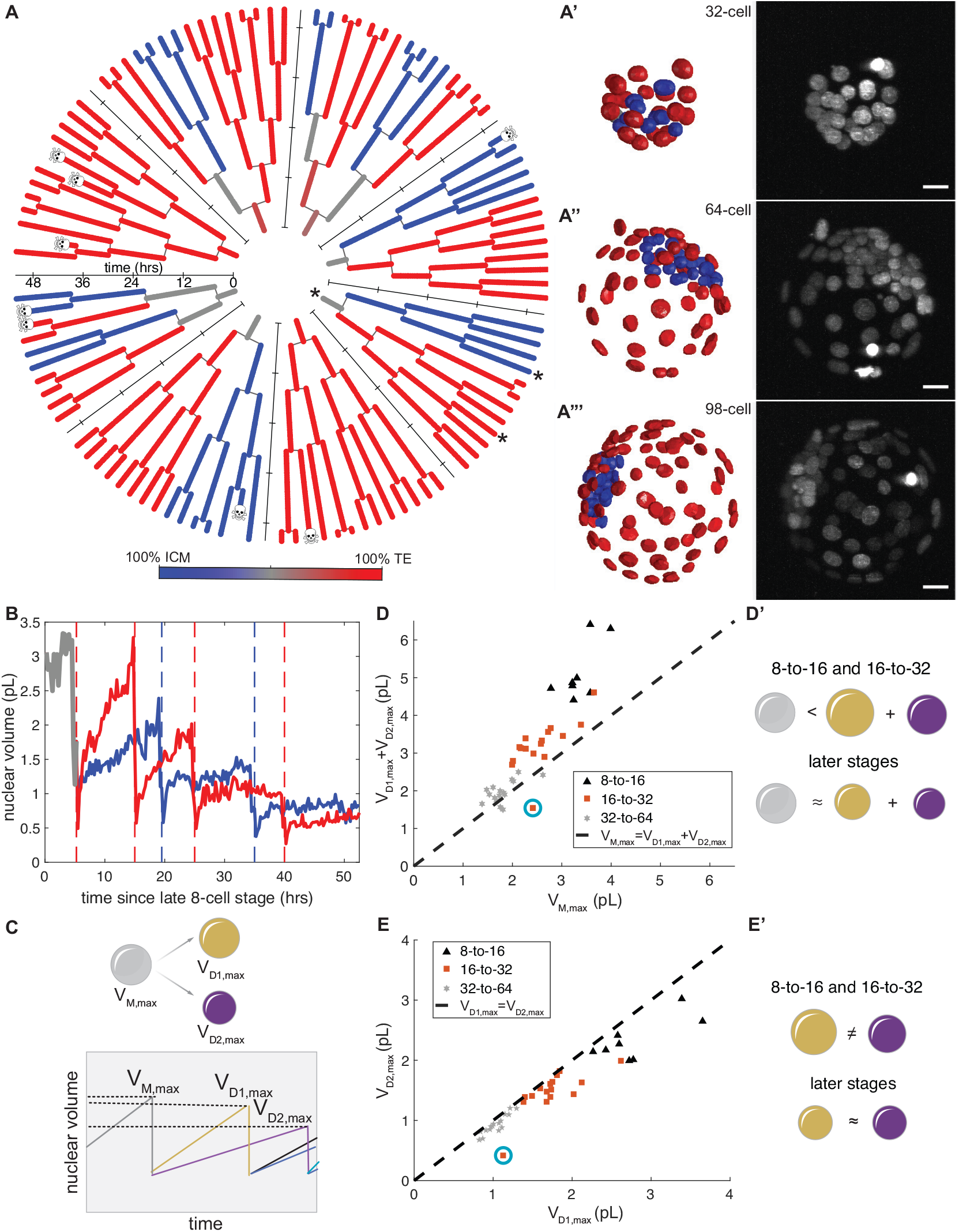
Probing nuclear dynamics from the 8-cell stage to the late blastocyst. (A) Lineage trees. Color indicates eventual contributions to the ICM and TE. Later times farther from center. Skull: cell death event. Paths between the asterisked root and two asterisked leaves in (B). (A’-A”‘) Max projections of H2B signal and segmentations, colored as in (A). Scale bars: 20 *μm*. (B) Nuclear volumes along the paths indicated by asterisks in (A). Trajectories are colored as in (A). Dashed lines: division events. (C) Cartoon of nuclear volume dynamics along the lineage tree. Gray, gold, and purple: mother nucleus, larger daughter nucleus, smaller daughter nucleus, respectively. (D,D’) Comparison of summed daughter nuclear volumes to that of the mother. (E,E’) Comparison of volumes of two daughter nuclei. In (D,E), cyan: outlier due to nuclear fragmentation.

**Fig 6.**
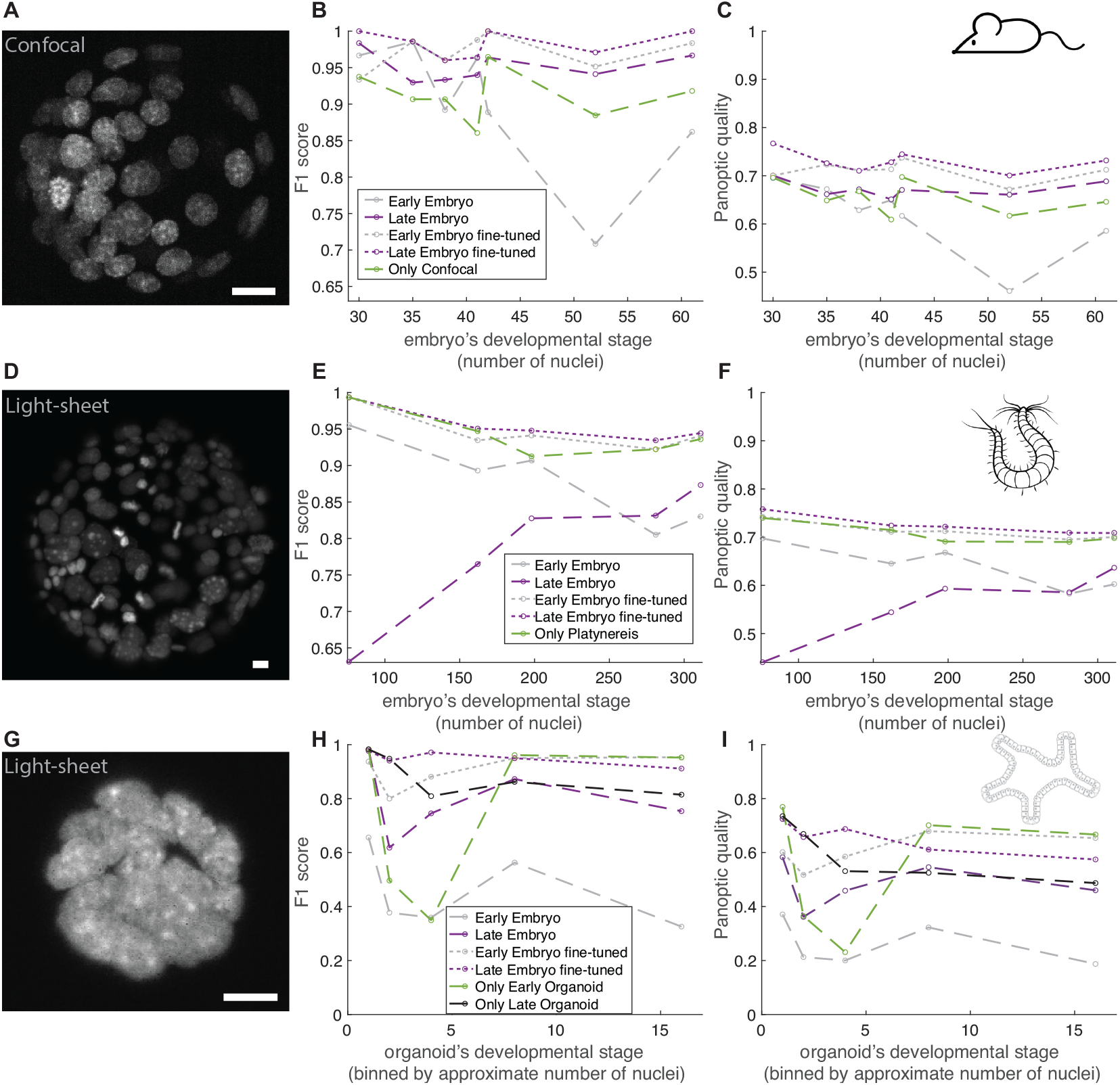
Generalization tests for different model systems and a different imaging modality. (A-C) For mouse embryos imaged via spinning-disk confocal microscopy, we generated ground-truth nuclear segmentation (see example max intensity projection in (C)). Panels in (A-B) indicate the performance of different models on the ground-truth test set. (D-F) Same as in (A-C) but for a ground-truth set of *Platynereis dumerilli* embryos [10]. (G-I) Same as in (A-C) but for a ground-truth set of intestinal organoids [9]. For each case, fine-tuning of our networks enables superior performance as compared to the network(s) trained on the system-specific ground-truth alone. All scale bars: 10 *μm*.

To track corresponding nuclear instances across time, we first corrected for movement of the embryo during image acquisition by computing a rigid transformation which aligns consecutive pairs of frames. Based only on nuclear centroids, we registered consecutive frames by utilizing the coherent point drift algorithm from [19]. Finally, we performed semi-automated lineage tracking on the registered nuclear instances. Our algorithm operates sequentially, matching nuclei to their predecessors in the previous frame. Non-dividing nuclei were tracked using nearest neighbor association between instance centroids. Additionally, for dividing nuclei, we used a heuristic based on the difference in nuclear volume between potential mother-daughter triples. For more details regarding segmentation correction, registration, and lineage construction, please see Description of Semi-automated Nuclear Tracking Methods in the methods section.

### 0.6 Using the analysis pipeline to generate lineage trees and characterize nuclear volumes and shapes from the 8-cell to late blastocyst stage

We used the pipeline (Figure 4) for the semi-automated analysis of an embryo from the 8-cell stage to the *≈* 100-cell stage (Figure 5(A-A”‘), S10 Fig). After *≈* 52 hours of development, by a combination of 98 division events and 8 death events, the embryo reached the 98-cell stage. At the final time point, we used nuclear position as a proxy to assign ICM and TE fates and found 27 ICM nuclei and 71 TE nuclei, in keeping with fate proportions previously reported [29]. To our knowledge, these are the longest lineage trees constructed for perimplantation development.

Next, we characterized changes in nuclear volumes and shapes in individual cell lineages in Figure 5(A) (see Quantifying nuclear shape properties in Methods). First, we examined how accurately our BlastoSPIM-trained models report on these features by comparing nuclear volume and aspect ratios in embryos with both ground truth and model-predicted segmentations. In comparing each model-predicted instance with its matched ground-truth instance, we found that the nuclear volumes and aspect ratios of the two matched well, particularly for the 8-cell stage up to the *≈* 80-cell stage (S5 Fig, S11 Fig). Therefore, for all nuclei throughout the time lapse in Figure 5(A), we calculated nuclear volumes and aspect ratios based on the corrected model predictions and treated those as a proxy for the ground-truth nuclear volumes and aspect ratios.

Since the nucleus’s volume relative to that of the cytoplasm (NC ratio) has been shown to impact cell cycles and gene expression in embryos of other species [30, 31], we first studied the dynamics of nuclear volumes from the 8-cell stage to the blastocyst stage. For preimplantation mouse embryos, the total embryo volume – excluding the cavity – is fixed, which means that cell divisions partition existing volume [32]. By contrast, analysis of fixed embryos has revealed that nuclear volumes do not downscale as dramatically as cell volumes [33]. To quantify in a live embryo how these increases in NC ratio arise, we analyzed nuclear volumes from generation to generation (*e.g*., from a mother at the 16-cell stage to 2 daughters at the 32-cell stage). Figure 5(B) shows an ICM and a TE example of nuclear volume trajectories from the root of a lineage tree (8-cell stage) to its leaves (*≈* 100-cell stage) (see S10 Fig). After each division, the nuclear volume grew approximately linearly and reached a peak immediately before the subsequent division (see Figure 5(C), S10 Fig). To compare nuclear volumes at different developmental stages in a way that is not affected by the asynchonous nature of cell cycle progression, we chose to use maximal nuclear volumes, which we measured for each cell immediately before mitosis.

We compared the maximal volumes of the daughter nuclei (*V*_*D*1,*max*_ and *V*_*D*2,*max*_) to that of their mother (*V*_*M,max*_) across developmental stages (Figure 5(C,D), S12 Fig(B)). If the nuclei were not down-scaling with developmental stage at all, then *V*_*D*1,*max*_ + *V*_*D*2,*max*_ would be expected to be *≈* 2*V*_*M,max*_. On the other hand, if the nuclei were down-scaling to fix the NC ratio, we would expect *V*_*D*1,*max*_ + *V*_*D*2,*max*_ to be *≈ V*_*M,max*_. For the 8-to-16 and 16-to-32 cell transitions, the sum of nuclear volumes of the two daughters (*V*_*D*1,*max*_ + *V*_*D*2,*max*_) was greater than *V*_*M,max*_, but less than *≈* 2*V*_*M,max*_. Thus, the daughter nuclei were indeed down-scaling, but were not simply halving the mother’s nuclear volume. By contrast, for the 32-to-64 cell stage, the sum of the daughter volumes approximately equaled that of the mother (Figure 5(C,D)). This suggests that nuclear growth is reduced at later developmental stages (see example trajectories in S10 Fig). Future work is required to uncover the specific biological mechanisms controlling developmental-stage-dependent nuclear growth.

During the 8-to-16 and 16-to-32 cell transitions, a previous study reported that differences in cell volume – particularly between ICM cells to TE cells – arise [21]. By comparing the volumes of daughter nuclei resulting from the same division (*V*_*D*1,*max*_ and *V*_*D*2,*max*_) across developmental stages (Figure 5(C,E), S12 Fig(C)), we tested when nuclear volume differences emerge in the embryo. For the 8-to-16 and 16-to-32 cell transitions, many of the divisions had pronounced asymmetries (Figure 5(E), S12 Fig(C)). By contrast, for the 32-to-64-cell transition, the daughter nuclei had approximately equal maximal nuclear volumes (Figure 5(C,E)). Since the nuclear volume asymmetries in earlier divisions often correspond to cases of two progeny of differing ICM/TE fate (S10 Fig), decreased asymmetries at the 32-to-64-cell transition may reflect the lack of mixed ICM/TE progeny in divisions after the 32-cell stage [7].

In addition to nuclear volumes, nuclear aspect ratios likely also depend strongly on developmental stage and ICM/TE cell fate. Since previous studies reported that at the 16-, 32-, and 64-cell stages, the TE cells have larger aspect ratios than ICM cells [21, 34], we asked whether and at what developmental stage TE nuclei develop higher aspect ratios than ICM nuclei. We found that TE nuclei developed high aspect ratios not during the 16-cell stage, but later during the 32-to-64 cell stages (S10 Fig,S12 Fig(G-H”‘)). This change in TE nuclear shape occurred highly asynchronously across the TE population (S10 Fig, S12 Fig(G-H”‘)) and sometimes occurred very quickly (S12 Fig(H”‘)). Additionally, although TE nuclei tended to have higher aspect ratios than ICM nuclei, the distributions significantly overlapped, and both the median TE and ICM nuclear aspect ratios increased by the *>*100-cell stage (S10 Fig). Interestingly, a subset of ICM nuclei experienced an increase in aspect ratio during the last 3-4 hours of the time-lapse (see S10 Fig(C’,D’)). This latter subset may correspond to primitive endoderm nuclei that flatten as the primitive endoderm forms a monolayer (Fig 1(B)) [35].

### 0.7 Generalization of our trained Stardist-3D models to different model systems and a different imaging modality

Since previous studies have reported that deep convolutional neural networks can generalize well to unseen datasets [5, 23, 36], we next tested whether our BlastoSPIM-trained models could generalize to other model systems and imaging modalities. In principle, our model’s performance on a different dataset could depend on the model system, the method for nuclear labeling, and the imaging modality. In each generalization test below, we first evaluated our early embryo and late blastocyst models on the test set without any additional training. Then, we updated the weights of our early embryo and late blastocyst models by training on a small set of ground truth data from the system of interest. We compared those so-called fine-tuned models to a model trained only on ground-truth data from the system of interest.

First, to test the generalization of BlastoSPIM-trained models on a different imaging modality, we generated a ground-truth set of 10 preimplantation mouse embryos from the *≈* 32-cell stage to the *≈* 64-cell stage on a spinning disk confocal microscope (see example in Figure 6(A)). We set aside 2 embryos for training and validation of a new “only confocal” model. We used that same set to update the weights of our two BlastoSPIM-trained Stardist-3D models. On the remaining embryos, the late blastocyst model – with and without training on confocal data – outperformed the “only confocal” model (Figure 6(B,C)). The fine-tuned early embryo model also outperformed the “only confocal” model. Thus, the late blastocyst model performed well out-of-the-box, and fine-tuning on a small training and validation set significantly improved the performance of both the early embryo model and the late blastocyst model. Thus, our models generalized well, alleviating the necessity of generating a large new ground-truth set of confocal data.

We quantified whether our models generalize to datasets from other model organisms. We evaluated our Stardist-3D models on a ground-truth set of live light-sheet images of *Platynereis dumerilli* embryos from the 38-to the 392-cell stage [10] (see example in Figure 6(D)), in which nuclei were labeled by microinjection of a fluorescent tracer. Applying our “early embryo” model to the five *P. dumerilli* images, we found that it performed well, at ⪆ 90% *F*_1_ score, on early *Platynereis* embryos, from the 76-to 198-cell stages. For later stages, the “late blastocyst” model outperformed the “early embryo” model by achieving an *F*_1_ score of *≈* 85% (Figure 6(E)). We, then, fine-tuned both of our Stardist-3D networks on a set of 4 *P. dumerilli* embryos. We used the same sets to train and validate a new network, which we call the “only platynereis” model. The fine-tuned “late blastocyst” model outperformed the “only platynereis” model (Figure 6(E,F)) across the test set. The superiority of the *F*_1_ score and the panoptic quality for the fine-tuned “late blastocyst” model was most pronounced for the *≈* 200-cell embryo. For the three test images at late stages (with 198, 281, and 311 nuclei), the fine-tuned network generated *≈* 30 fewer errors, including false positives and false negatives, and increased the mean IoU for matched instances at each timepoint. Notably, our models generalized to images of *P. dumerilli* embryos despite several differences in this data compared to images of mouse embryos, such as highly variable nuclear sizes and different nuclear textures (dense heterochromatic foci).

Finally, we tested our models’ generalizability to another system, intestinal organoids [9] (Figure 6(G)). While our “early embryo” and “late blastocyst” models did not perform as accurately out-of-the-box as they did on the previous two sets, fine-tuning of both of these models on sets of images of 10 organoids with ground-truth significantly improved their performance (Figure 6(H,I)). For example, the fine-tuning of the early embryo model on early organoid data (*<* 14 nuclei) and of the late blastocyst model on late organoid data (*≥* 14 nuclei) allowed these to outperform the “only organoids” models (trained on early and late organoid data, respectively). Surprisingly, the best improvement in performance relative to the “only organoid” models occurred at the 2-4 cell stages (Figure 6(H,I)). At these stages, a lumen has not yet formed or is small, and the cells’ large nuclei are very closely juxtaposed and flattened. The large training set of blastocyst embryos (in BlastoSPIM 1.0 and 2.0) enabled our models to avoid merging of these nuclei in the segmentation.

Thus, our BlastoSPIM-trained models, either ”out-of-the-box” or fine-tuned with minimal ground truth from another system, can greatly improve nuclear segmentation accuracy in different types of imaging data (Figure 6).

## Conclusion

To understand how individual cells’ behaviors contribute to morphogenetic events, biologists acquire staggering amounts of time-lapse images of these processes. Quantifying the properties and behaviors of individual cells in such image series requires instance segmentation: identifying which voxels belong to which object. Although many measurements require segmentation of entire cells, instance segmentation of nuclei is useful for estimating the relative positions of cells, classifying by mitotic stage, and measuring the expression of nuclear-localized factors. Nuclear instance segmentation is challenging for several reasons, including nucleus-to-nucleus proximity, variations in nuclear shape, voxel anisotropy, and low SNR. Since the application of supervised machine learning methods to instance segmentation often requires relatively large ground-truth datasets, here we generated a publicly available ground-truth dataset, called BlastoSPIM, which is the largest 3D dataset of nuclear instance segmentation ground truth of its kind. Such large, fully annotated datasets are an extremely useful resource, including for the benchmarking of different methods. Here, by comparative analysis of seven different neural networks on this new dataset, we have shown which of these networks best addresses the challenges of nuclear segmentation in the preimplantation mouse embryo (Figure 2).

Our comparative analysis revealed state-of-the-art performance by Stardist-3D (early embryo model) across developmental stages. From the 8-cell stage up to the 64-cell stage, Stardist-3D’s *F*_1_ score remained above 95 %, and its panoptic quality at *≈* 80 % (Figure 2(A)). In contrast, the performance of other methods varied, with Cellpose and RDCNet producing many false positives particularly at early developmental stages, and U3D-BCD and UNETR merging several nuclei for the 64-cell stage and later stages (Figure 2). To further improve segmentation performance at later stages of preimplantation development, we hand-annotated a second ground truth dataset of nuclei in late blastocyst embryos and trained a second Stardist-3D model (late blastocyst model), of which the *F*_1_ score remained above 90 % for the *>* 100-cell stage embryos (Figure 3(A)). Therefore we not only present trained Stardist-3D models with superior performance for nuclear instance segmentation in time-lapse images of early mouse embryos, but share large ground truth datasets (BlastoSPIM 1.0 and 2.0), which will be an important resource for evaluating the performance of future CNN architectures because of the dataset’s size and quality and nuclear density relative to other currently available datasets (S2 Table).

Due to our interest in studying preimplantation mouse development in live images, we integrated our Stardist-3D models (the early embryo and late blastocyst models) into a complete image analysis pipeline (Figure 4), including time-lapse acquisition, instance segmentation, segmentation correction, nuclear centroid registration and lineage tracking. The models and the code underlying the pipeline are made publicly available at blastospim.flatironinstitute.org. We used this pipeline to analyze a time-lapse image from the 8-cell stage to the *≈* 100-cell stage. These segmentations revealed oscillations of nuclear volume with the cell cycle: volumes gradually increased throughout interphase and peaked just before mitosis, resulting in a sudden volume drop. We probed how nuclear volumes downscale from earlier developmental stages to later stages. Whereas previous studies have quantified this effect based on fixed embryos or live images without tracking, we used our lineage trees to compare the max volume of each mother nucleus to the max volumes of the resulting daughter nuclei (Figure 5(C,D), S12 Fig(B)). We found that nuclei did indeed downscale in volume over time, but that the max nuclear volumes of the daughter were not simply half that of the mother (see S12 Fig for comparison to ground-truth examples). Additionally, sibling nuclei often differed significantly in nuclear volume at early developmental stages (Figure 5(C,E), S12 Fig(C)), but such asymmetries decreased considerably by the 32-to-64 cell transition (Figure 5(C,E)). We expect that our instance segmentation models will enable many more insights into the development of preimplantation mouse embryos, including into the fate decision occurring within the ICM.

The usefulness of datasets like BlastoSPIM often extends to images acquired by different modalities or of different model systems. Here we tested whether: (1) models trained on BlastoSPIM can be applied off-the-shelf to segment nuclei in other contexts and (2) BlastoSPIM can be used to improve accuracy in segmentation via pre-training when limited ground-truth data is available. Our generalization tests extended across three datasets: preimplantation mouse embryos imaged by spinning-disk confocal microscopy, intestinal organoids imaged by light-sheet microscopy [9] and *Platynereis dumerilli* embryos imaged by light-sheet microscopy [10]. Our early embryo model and late blastocyst models worked well out-of-the-box for the spinning-disk confocal set and up to the 200-cell stage for the *Platynereis dumerilli* embryos. Furthermore, fine-tuning of our models on the spinning-disk confocal set and the *Platynereis dumerilli* embryos improved model performance, both in terms of *F*_1_ score and panoptic quality, relative to a model trained on system-specific data alone. Finally, fine-tuning of our models on the intestinal organoids set improved performance relative to an “organoid only” model, particularly for stages when the nuclei were densely packed without a separating lumen. Thus, just as with other large ground-truth datasets (for example, ImageNet for object recognition in 2D images), finetuning or transfer learning from models trained on BlastoSPIM should improve performance on related tasks.

The generalizability of our model fills a clear need since many publicly available models work only in 2D, segment only cell boundaries, or are trained only on high SNR images [37]. Given our model’s performance even without fine-tuning, small hand-corrections of our model’s predictions on a different biological system could be used to generate training data, as long as that system’s nuclei satisfy the star-convexity assumption of Stardist. We expect that BlastoSPIM and our Stardist-3D models, in conjunction with other publicly available datasets and pre-trained models [38], will play a key role in the development of truly generalist models. Furthermore, the integration of BlastoSPIM-trained models into a larger analysis pipeline enabled the construction of lineage trees, which revealed the temporal dynamics of individual nuclei as fate decisions transpire. This work is thus a crucial step towards fully automated (3+t)-D analysis of early mouse development, and the full pipeline will likely prove useful for the analysis of developmental dynamics of other model organisms.

## Supplementary Information

**S1 Table.**
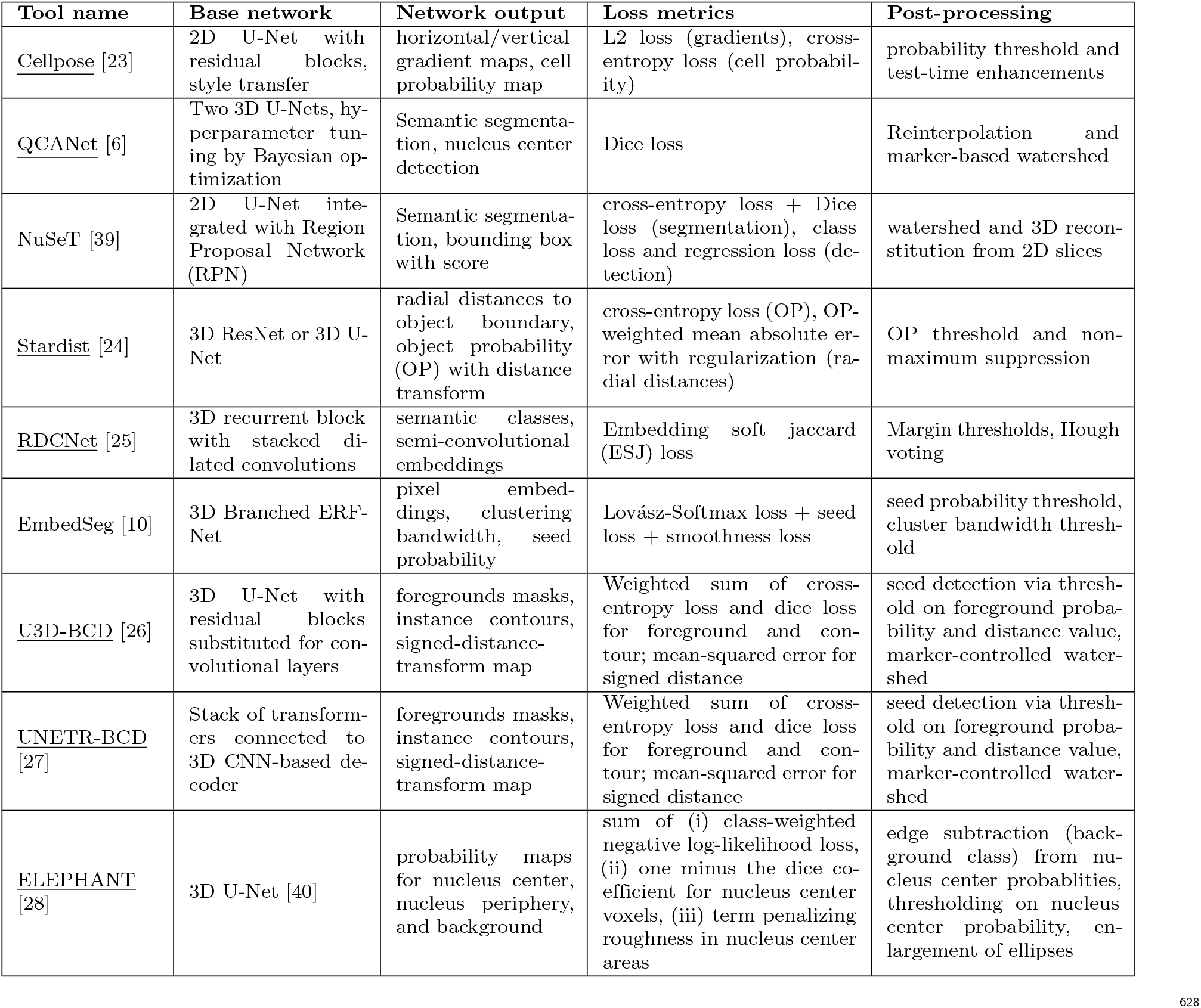
Tools for 3D nucleus segmentation. Underlined: methods used for benchmarking here

**S2 Table.** Ground-truth, three-dimensional annotations of nuclei. Other examples of publicly available ground-truth data sets for instance segmentation of nuclei. *The entire ground-truth dataset used in [6] contains more than 6000 time-series of early mouse embryos, of which only 165 have been made publicly available.

| Name | Microscopy | Nuclear Labeling | Sample | Image Count | Network results |
| --- | --- | --- | --- | --- | --- |
| NucMM-Z [26] | Serial-section electron microscopy | N/A | Zebrafish brain | 1 | Cellpose3D, Stardist-3D, U3D-BCD |
| NucMM-M [26] | Micro-CT | N/A | Mouse visual cortex | 1 | Cellpose3D, Stardist-3D, U3D-BCD |
| BBBC050* [6] | Confocal microscopy | H2B-mRFP1, H2B-mCherry | Pre-implantation mouse embryo from the pro-nuclear stage to 53-cell stage | 165 | QCANet, 3D U-Net, 3D Mask R-CNN |
| *C. elegans* developing embryo [41] | Confocal microscopy | histone-GFP | *C. elegans* embryo between 2-cell stage and ≈300-cell stage | 9 | QCANet, 3D U-Net, 3D Mask R-CNN |
| *Platynereis*-Nuclei-CBG [10] | Light-Sheet Microscopy | Fluorescent nuclear tracer injected | *Platynereis dumerilii* embryo between 0 and 16 hours post-fertilization | 9 | Cellpose3D, Stardist-3D, EmbedSeg |
| *Platynereis*-ISH-Nuclei-CBG [10] | Confocal Microscopy | DAPI | *Platynereis dumerilii* specimens 16 hours post-fertilization | 2 | Cellpose3D, Stardist-3D, EmbedSeg |
| *Parhyale hawaiiensis*-Nuclei [24] | Confocal Microscopy | H2B-eGFP | *Parhyale hawaiiensis* embryo between 46 hours post-amputation (hpa) and 110 hpa | 6 | U-Net, Stardist-3D, Cellpose3D, EmbedSeg |
| *C. elegans*-Nuclei [24] | Confocal Microscopy | DAPI | *C. elegans* embryo at the 558-cell stage | 28 | U-Net, Stardist-3D |
| Mouse-Skull-Nuclei-CBG [10] | Confocal Microscopy | DAPI | Nuclei from the skull of developing mouse embryos | 2 | Cellpose3D, Stardist-3D, EmbedSeg |
| Peri-implantation mouse embryos [42] | Confocal Microscopy | Antibody staining | Peri-implantation mouse embryos | 35 | 3D U-Net |

**S3 Table.** Segmentation of H2B signal in mitotic cells. For mitotic nuclei, there were only 3 late-stage images (2 at the 64-cell stage, and 1 at 100-cell stage) with 6 examples. Nuclei were considered mitotic if they were in metaphase or anaphase. For these late-stage embryos, there are many small nuclei tightly packed together. All the methods missed at least one mitotic nuclei at IoU=0.5 with UNETR-BCD performing the best with only one.

| Method name | Average IoU | Misses (IoU < 0.1) | Misses (IoU < 0.5) | FP (IoU = 0.5) |
| --- | --- | --- | --- | --- |
| Cellpose | 0.52 | 1 | 2 | 2 |
| QCANet | 0.27 | 1 | 5 | 5 |
| Stardist | 0.46 | 1 | 2 | 2 |
| RDCNet | 0.37 | 0 | 4 | 4 |
| U3D-BCD | 0.41 | 1 | 4 | 4 |
| UNETR-BCD | 0.65 | 0 | 1 | 1 |
| ELEPHANT | 0.37 | 0 | 4 | 4 |

**S4 Table.** Segmentation of Polar Bodies. There were 13 images in which a total of 15 polar bodies were labelled as such. All the methods detected all the polar bodies (except one that Cellpose missed at IoU=0.1) but with varying degrees of success in how well they were segmented. At IoU=0.5, most methods missed a few, except for RDCNet which was able to detect them all. We hypothesize that RDCNet which uses much fewer parameters given its recursive design, is better able to learn to detect polar bodies using the few examples in the training set.

| Method name | Average IoU | Misses (IoU < 0.1) | Misses (IoU < 0.5) | FP (IoU = 0.5) |
| --- | --- | --- | --- | --- |
| Cellpose | 0.47 | 1 | 7 | 6 |
| QCANet | 0.35 | 0 | 14 | 14 |
| Stardist | 0.66 | 0 | 2 | 2 |
| RDCNet | 0.79 | 0 | 0 | 0 |
| U3D-BCD | 0.66 | 0 | 4 | 4 |
| UNETR-BCD | 0.70 | 0 | 2 | 2 |
| ELEPHANT | 0.60 | 0 | 5 | 5 |

**S1 Fig. Segmentation tasks applied to images of a pastoral scene [43] and of a mouse embryo**. (A) Raw image to be segmented. From top to bottom, an image of cows in a pasture, maximum intensity projections of 3D image of 16-cell mouse embryo, a z-slice from the 3D image. 3D image has dimensions (83.4 *μm*, 83.4 *μm*, 68 *μm*). Scale bar: 10 *μm*. (B) Semantic segmentation for images in (A). (C) Object detection for images in (A). (D) Instance segmentation for images in (A).

**S2 Fig. SNR of each image in the BlastoSPIM dataset**. (A-B) Histogram, for annotated images in the original BlastoSPIM set and in the corrected late blastocyst segmentations, respectively, of the difference between mean foreground intensity and the mean background intensity. Black line: Cutoff for separating low SNR from moderate-to-high SNR in the original BlastoSPIM dataset.

**S3 Fig. *F*_1_ score and panoptic quality across IoU thresholds for seven methods. Analogous to Figure 2(A,B).** (A,A’) *F*_1_ score and panoptic quality, respectively, for an IoU cutoff of 0.2. (B,B’) Same as in (A,A’), for an IoU cutoff of 0.3. (C,C’) Same as in (A,A’), for an IoU cutoff of 0.4. (D,D’) Same as in (A,A’), for an IoU cutoff of 0.6. Note that Figure 2(A,B) are based on an IoU cutoff of 0.5.

**S4 Fig. *F*_1_ score and panoptic quality across IoU thresholds for early embryo, late blastocyst, and all stages models. Analogous to Figure 3(A,B)**. (A,A’) *F*_1_ score and panoptic quality, respectively, for an IoU cutoff of 0.2. (B,B’) Same as in (A,A’), for an IoU cutoff of 0.3. (C,C’) Same as in (A,A’), for an IoU cutoff of 0.4. (D,D’) Same as in (A,A’), for an IoU cutoff of 0.6. Note that Figure 3(A,B) are based on an IoU cutoff of 0.5.

**S5 Fig. Comparing ground-truth and model-predicted nuclear volumes across embryonic stages**. For the combined ground-truth set used in Figure 3(A,B), if a ground-truth instance is matched to a model-predicted instance by an IoU of at least 0.5, we plot the ground-truth nuclear volume against the predicted nuclear volume. Each panel (A-G) is labelled with the corresponding developmental stage. See legend for meaning of dashed lines. Note that the large ground-truth instances at the 8-cell stage correspond to a couple of ground-truth images in which an annotator overestimated nuclear sizes because of low SNR. Encouragingly, our model predicted volumes for these instances which are closer to range of the rest of ground-truth volumes. For the 8-cell stage, the second *R*^2^ value is for the set with the overestimated ground-truth volumes removed.

**S6 Fig. Nearest-neighbor distances between nuclei decrease dramatically during development**. (A-D) Example z-slices and quantification for 16-cell (A), 32-cell (B), 50-to-64-cell (C), and *>*90-cell (D) embryos. The first two rows contain images and corresponding annotations. Each red arrow indicates the nucleus’s effective radius, the radius of a sphere of equivalent volume. The gray lines indicate examples of shortest surface-to-surface distance. The third and fourth rows show that the effective radius and the shortest surface-to-surface distance decrease during development. Illustrations in the bottom histograms show that the latter decreases more than the former. Median of histogram in black. Scale bar: 10 *μm*.

**S7 Fig. Quantifying how well star-convex approximation applies to nuclear shapes in ground-truth time series data**. We fit each nucleus to a star-convex shape, using 128 rays. For a single embryo, for which we have annotated ground truth for 89 consecutive timepoints (time points acquired every 15 minutes), we plot a box for each time to illustrate how well this fit performs, in terms of IoU. When all nuclei are in interphase, the star-convex fit performs quite well, at more than 90 percent IoU between the ground truth and the model-generated instance. During the transition from the 16-cell stage to the 32-cell stage and from the 32-cell stage to the 64-cell stage, the fit quality degrades. A small number of nuclei, about five in this time series cannot be fit by a star-convex shape, resulting in an IoU of less than 40 percent. We expect that the outlier nuclei (red) – which are not well fit by a star-convex shape – are likely mitotic, most likely in either metaphase or anaphase when the shape of the condensed chromatin is often complex. Black dashed line: the number of nuclei versus time.

**S8 Fig. Qualitative evaluation of failure modes for late blastocyst model on images of early embryos, particularly those with low SNR**. Blue: Model prediction which is a true positive. Green: Model prediction which is a false positive. (A) z-slice containing only a polar body. Note that the late blastocyst model predicts a couple of false positives around the polar body. (B) z-slice in which debris and a reflection from the image chamber are falsely segmented into instances. (C) z-slice in which a false positive is detected away from the embryo (see region with true positives).

**S9 Fig. Quantitative and qualitative evaluation of failure modes for the early embryo model and the late blastocyst model on images with particularly low SNR**. (A,B) F1-score and panoptic quality, respectively, for the low SNR test set, binned by developmental stage. (C,D) Representative xy slices from an *≈*90-cell embryo and an *≈*64-cell embryo, respectively. Results from the early embryo model in (C) illustrate its tendency to improperly merge closely juxtaposed nuclei into a single instance. Results from the late blastocyst model in (C,D) illustrate its tendency to produce false positives, sometimes in regions with nuclei which are blurred or haloed due to imaging artifacts. It is worth noting that these images, particularly (D), have extraordinarily low SNR, which makes even manual annotation difficult. The results in this supplemental figure, thus, represent some of the most difficult test images to segment.

**S10 Fig. Dynamics of nuclear aspect ratios and nuclear volumes from Figure 5(A)**. (A,A’) Nuclear volume and nuclear aspect ratio, respectively, for two paths in the lineage tree from the same root. When the paths split at a division, one resulting daughter’s line is dashed, while the other’s remains solid. Vertical dashed lines indicate division events. (B,B’) Nuclear volume and nuclear aspect ratio, respectively, for two paths illustrated in Figure 5(B). Colors as in the tree in Figure 5(A). (C,C’) Nuclear volume and nuclear aspect ratio, respectively, for two paths from the same root to leaves with different fates (one ICM, one TE). Arrow indicates increased aspect ratio of ICM nucleus towards the end of the time lapse. (D,D’) Same as in (C,C’), but for two different paths.

**S11 Fig. Comparing ground-truth and model-predicted aspect ratios across embryonic stages**. For the combined ground-truth set used in Figure 3(A,B), if a ground-truth instance is matched to a model-predicted instance by an IoU of at least 0.5, we plot the ground-truth nuclear aspect ratio against the predicted nuclear aspect ratio. Each panel (A-G) is labelled with the corresponding developmental stage. See legend for meaning of dashed lines.

**S12 Fig. Dynamics of nuclear volume and aspect ratios in 2 ground-truth lineages**. (A,A’) Two fully ground-truth lineages, including both ground-truth nuclear annotation and nuclear tracking. The tree coloring indicates ICM-TE lineage contribution as in Figure 5(A). (B) Plot analogous to Figure 5(D), for the two ground-truth lineages (one in cyan, one in red). Each point represents one mother nucleus from the 16-cell stage giving rise to two daughter nuclei at the 32-cell stage. (C) Plot analogous to Figure 5(E), for the two ground-truth lineages (one in cyan, one in red). (D) During the 32-cell stage (see the horizontal black line), the TE nuclear volumes become statistically significantly bigger than the ICM nuclear volumes (comparison based on rank sum test at each time). Dashed lines: mean nuclear volumes for the ICM and TE nuclei in (A). Solid lines: result of fitting lines to inter-division nuclear trajectories, then averaging those lines for the ICM and TE separately. (E) Same as in (D) but for the lineage in (A’). (F) Cartoon summarizing nuclear volume dynamics in the lineages. By fitting a line to each inter-division interval at the 32-cell stage, we extract the value of the linear fit immediately after the division (horizontal dashed lines), the growth rate (indicated by solid black lines), and the time of division (vertical dashed line). (F’, F”, F”‘) For each of the ground truth trees, statistical comparisons between ICM and TE linear fits. p-values result from rank sum test. (G-G”‘) Example nuclear aspect ratios for the lineage in (A) Each segment is colored as in (A). If two daughters are of same fate, one is plotted as a dashed line. Lines end when next division occurs. (H-H”‘) Same as (G-G”‘) but for the lineage in (A’) Black arrow indicates sudden change in nuclear aspect ratio.

### S1 File. Sequence file for H2B-miRFP720 mouse line

#### Network Implementation Details

Each model was trained with data from 482 3D images of whole embryos. Each embryo was cropped into 8 to 16 patches depending on the size for a total of 4363 patches. Each patch had a resolution of 64×256×256. The patches overlap such that all voxels of a nucleus were fully contained in at least one patch. The raw intensity images were bit-shifted by four bits to the right, so that all voxel intensities are in the range between 0 and 255. Any value still above 255 was capped at 255.

#### RDC Net

For all hyperparameter combinations sampled for training, a few were held constant. The down sampling factors were chosen to be 1, 10, and 10, for the z, x, and y directions, respectively, to account for anisotropy. Spatial dropout was chosen to be 0.1, following the original paper. All networks were trained for a maximum of 200 epochs, batch size of 2, with the Adam optimizer and Cosine Decay Restarts scheduler with learning rates from 10-3 to 0. The set of model weights that resulted in the lowest validation loss across all epochs was saved. The patch size was either the original crop (64×256×256) or 32×256×256 (32 random, consecutive Z slices from the original crop). The number of groups (parallel stacked, dilated convolution blocks with shared weights), dilation rates, number of channels per group, number of iterations, and the margin parameter were also adjusted to observe their effects on network performance.

During inference on test images, each raw image was broken into patches with the same size as those the model was trained on. The test patches were passed through the model and the resulting label patches were stitched together by discarding redundant masks and any masks touching the patch boundaries, assuming each nucleus is located at the center of at least one patch.

#### Cellpose

Cellpose is called a generalist method for cell and nuclei instance segmentation. It is based on a 2D U-Net with residual blocks and style transfer. The objects are modeled as a diffusion gradient. The output is composed of horizontal and vertical gradient maps and a segmentation probability map. Since the original cellpose model is 2D, the 3D patches were broken into 2D slices for training. The source code was modified to include a data loader, since the size of the 2D training set is orders of magnitude larger than the original Cellpose dataset. Models were trained for a maximum of 1000 epochs, either from scratch or from a pretrained Cellpose model. Test images were down-sampled by a factor of 0.5 in X and Y to improve performance since Cellpose is prone to over-segmentation for our full-resolution images in 3D. The patching and stitching method was the same as for RDCNet.

#### Stardist

3D Stardist was trained with patches of 32×256×256 sampled from the full size patches. Input intensity was normalized capping values below 1% and above 99.8%. For sampling the star convex in 3D, we used 96 rays with a grid of 1×4×4 to compensate for the anisotropy. Data augmentation included 2D flips and grid warping.

#### U3D BCD and UNETR

In U3D BCD, a 3D Residual U-Net, and UNETR, which uses a transformer as an encoder, the instance segmentation problem is broken down into learning hybrid representations i.e., semantic, contour and signed distance transform maps with the help of neural networks, and using watershed algorithm to separate instances.

UNETR encoder’s transformer uses an embedding dimension of 768, the input volume is patched into volumetric tokens of dimensions 16 ×16×16, and multi-head self-attention is performed with 12 heads. Augmentations, in the form of randomized brightness and contrast, flips, rotations and elastic deformations, were used. Finally, the input volumes were randomly cropped to 16×128×128, before passing them through the network. Adam optimizer with decaying learning rate was chosen for training. Weighted sum of Binary Cross Entropy (BCE) Loss and Dice Loss is taken for foreground and contour masks, while Mean Squared Error (MSE) was utilized for signed distance transform map predictions. Inference is performed by processing overlapping sliding windows across the large volumes of testing set. During post-processing, the multi-channel outputs from networks are combined by thresholding them appropriately to find instance seeds (or markers). Similarly, a more relaxed threshold on the outputs is used to obtain the foreground mask. Thereafter, marker-controlled watershed algorithm can be used with the help of seeds and predicted distance map to find instances.

#### QCANet

For QCANet, the original ground-truth set (BlastoSPIM 1.0) had to be converted to a set of nuclear centroids and semantic segmentation. The nuclear centroid image was computed by first down-sampling the xy-resolution by 4 and then by setting the value any voxels within 2.5 *μm* of a nuclear centroid to 1. The semantic segmentation was computed by setting all labelled regions in the instance segmentation to 1 and downsampling by 4 (using scikit-image: block reduce based on maxima). The instance segmentation was down-sampled in the same way, for comparing model predictions to ground truth. In all cases, the down-sampled ground-truth always had the same dimensions: 64×173×169.

Images were read in as unsigned 16-bit tiffs, with the z-resolution about 2.5 times less than the xy-resolution (as done in the original QCA study). To handle the very bright polar bodies in our images, we changed QCANet’s image normalization. Input intensity was normalized capping values below 1% and above 99.8% and then normalized using CSBDeep’s normalize function. Both the training and validation sizes for both nsn and ndn networks were 5, with augmentation. For all training, the epoch number was set to 200.

#### ELEPHANT

To train a 3D ELEPHANT model, we converted our ground truth data into ellipsoid labels, using the script generate seg labels.py in their latest (and continuously updated) main ELEPHANT repository (https://github.com/elephant-track/elephant-server). In the label generation step, the center ratio was set to 0.7. In the training step, the script train.py was used. We trained a model extending the pre-trained versatile model with our dataset of 64×256×256 crops using an additional crop size of 24×192×192, a batch size of 10, and for 40 epochs. We used the class weights for the negative log-likelihood loss (*class*_*weights*_): nucleus center=200, nucleus periphery=100, background=1, with a learning rate (lr) of 0.005. The validation dataset was built as a subset of the dataset by picking up every 10 image/label pairs. The final model was selected based on the validation set and was from the last epoch (i.e., epoch 40). The training was done in approximately 7.6 hours. In the inference step, the script elephant-detection.py was used with the following parameters: scales of 2.0×0.208×0.208 μm, a patch size of 24×256×256, a batch size of 10, a minimum radius (*r*_*min*_) of 2 μm, a maximum radius (*r*_*max*_) of 10 *μm*, a center ratio (*c*_*ratio*_) of 0.7, and a probability threshold (*p*_*thresh*_) of 0.5.

#### Training with Synthetic Data

To counter the limited number of samples with densely-packed nuclei, we generate artificial samples to learn generalized features. This is made possible by modeling nuclei as 3-dimensional Gaussian kernels, of dimensions x, y, z where x, y *∈* [100, 150] and z *∈* [3, 6]. Elastic deformations, randomized lighting, and addition of noise are done to match SNR ratios with that of actual data-set. The models are pre-trained with this simulated data, allowing the network to fine-tune its predictions on the actual data-set.

### Model Generalization Tests

#### Mouse blastocysts image by spinning disk-confocal

The generation of raw images and corresponding ground truth annotation is described in the Methods. The voxel resolution for these images is the same as that of our BlastoSPIM datasets. The validation set for the fine-tuned models and for the “only confocal” model is a single embryo with 50 nuclei, in 32×256×256 patches. The training set for the fine-tuned models and for the “only confocal” model was is a single embryo with 58 nuclei, in 32×256×256 patches. All models were trained as described in the Stardist section of Network Implementation Details. The fine-tuned models and for the “only confocal” model were trained in the same way, with the only difference being in the weights of the network at the start of training. Further improvement of the fine-tuned model relative to the “only confocal” model could possibly be obtained by lowering the learning rate or fixing some network weights during fine-tuning.

For the “early embryo” model and “late blastocyst” models results without fine-tuning, we combined the validation and training embryos described above (one embryo with 50 nuclei and another with 58 nuclei) to optimize only the nms threshold and probability threshold (using the built-in optimize thresholds) without tuning any weights in the network.

#### *Platynereis dumerilli* embryos

The raw images and corresponding ground truth annotation for *Platynereis dumerilli* embryos are from [10]. The voxel resolution for these images is 0.406 *μm* in xy as compared to 0.208 *μm* for the BlastoSPIM datasets, while the z resolutions of both sets are equal. We, thus, upsampled the images of *Platynereis dumerilli* embryos by a factor of 2 in xy (by interpolation with rescale from sci-kit image). The validation set for the fine-tuned models and for the “only *Platynereis*” model contains half of 32×256×256 patches of 2 images, one with 38 nuclei and the other with 392 nuclei. The rest of the 32×256×256 patches of these 2 images, in addition to 32×256×256 patches of images with 113 nuclei and 261 nuclei, are in the training set.

All models were trained as described in the Stardist section of Network Implementation Details. The fine-tuned models and for the “only confocal” model were trained in the same way, with the only difference being in the weights of the network at the start of training. Further improvement of the fine-tuned model relative to the “only confocal” model could possibly be obtained by lowering the learning rate or fixing some network weights during fine-tuning.

For the “early embryo” model and “late blastocyst” models results without fine-tuning, we combined the validation and training embryos described above to optimize the nms threshold and probability threshold without tuning any weights in the network.

#### Intestinal Organoids

The organoids dataset was provided by the Liberali Lab (see [9]). It contained 3 sequences of live intestinal organoids grown from single cells, 2 that were budding and 1 with a growing enterocyst. As compared to nuclear shapes in early preimplantation embryos, the nuclear shapes in intestinal organoids vary considerable across time and across different cells. Additionally, in images of intestinal organoids, the large nuclei are closely juxtaposed even before the formation of a lumen. We used frames 21 to 350 of the first budding organoid (starting with 2 nuclei and reaching *≈* 40 nuclei) for training and frames 21 to 350 of the enterocyst (starting with 1 nucleus and reaching *≈* 30 nuclei) for testing. We used these frames because these images had label images which appeared to be the most accurate with respect to the raw data.

We were given the original raw images and the deconvolved images but found that the models performed better with the raw images. This is not surprising, since the BlastoSPIM-trained models were trained on raw images as well. Denoising and deconvolving did not improve network performance although they improve the visual appearance for human viewing. Images were acquired using light-sheet recordings performed with a dual-illumination inverted light-sheet microscope. Images were acquired with an x-y resolution of 0.26 microns and 2.0 *μm* between slices.

### Computational Resource Requirements

For training of any of the models presented in this study, either a 40 GB A100 or a 32GB V100 was used. Training was not parallelized across multiple GPUs. Although the use of a GPU tends to speed up the process of model inference, a GPU is not required for that step. For inference, we requested 150GB memory on nodes with 2.6 GHz Intel Skylake CPUs. With 16 threads requested it takes on average 120-130s per image. Additionally, for the use of our models on new data from different model systems or imaging modalities, the optimization of the nms threshold and probability threshold – as opposed to the model weights themselves – with the use of the Stardist-3D function optimize thresholds does not require a GPU.

## Author contributions

Conceptualization, E.P., S.S., H.N., L.B., B.S.; Data curation, H.N., A.W., A.B., R.K., Z.G.; Formal analysis, H.N., B.S., L.B., P.G., J.S., D.D., M.A.; Funding acquisition, E.P., R.K.; Investigation, R.K., B.J., A.K., M.C., H.N., B.S., D.D., M.A., P.G., J.S., L.B.; Methodology, B.S., L.B., P.G., J.S., H.N., E.P., B.J., D.D., M.A., A.K., A.B.; Project administration, L.B., E.P., S.S.; Resources, B.J., A.K., Z.G.; Software, H.N., B.S., P.G., J.S., A.B., A.W., L.B., D.D., M.A.; Supervision, E.P., L.B., S.S.; Validation, H.N., A.B., L.B., D.D., R.K., M.C.; Visualization, A.W., H.N., D.D.; Writing – original draft, H.N., L.B., B.S., E.P.; Writing – review and editing, S.S., E.P., L.B., H.N., D.D., M.A.

## Acknowledgments

We thank Lucy Reading-Ikkanda (Flatiron Institute) for figure artwork and Marta Baziuk (Princeton), Alana Bernys (Princeton) and Christopher Catalano (Princeton) for work on ground-truth annotations. We thank Prisca Liberali and Gustavo Quintas Glasner de Medeiros (Friedrich Miescher Institute for Biomedical Research) for the intestinal organoids data. We thank Ko Sugawara (Ecole Normale Superieure de Lyon) for assistance with training ELEPHANT. This publication was made possible by grant numbers R01HD110577 and R01HD107026 (Posfai) from the National Institutes of Health. Research reported in this publication was supported by the National Center for Advancing Translational Sciences (NCATS), a component of the National Institutes of Health (NIH) under award number TL1TR003019. The content is solely the responsibility of the authors and does not necessarily represent the official views of the National Institutes of Health.

## References

1. X. Lou, M. Kang, P. Xenopoulos, S. Muñoz-Descalzo, A.-K. Hadjantonakis, A rapid and efficient 2D/3D nuclear segmentation method for analysis of early mouse embryo and stem cell image data, Stem Cell Reports 2 (3) (2014) 382–397. doi:10.1016/j.stemcr.2014.01.010.

2. D. J. Barry, C. Gerri, D. M. Bell, R. D’Antuono, K. K. Niakan, GIANI – open-source software for automated analysis of 3D microscopy images, Journal of Cell Science 135 (10) (2022) jcs259511. doi:10.1242/jcs.259511. URL https://journals.biologists.com/jcs/article/135/10/jcs259511/275435/GIANI-open-source-software-for-automated-analysis

3. G. Blin, D. Sadurska, R. Portero Migueles, N. Chen, J. A. Watson, S. Lowell, Nessys: A new set of tools for the automated detection of nuclei within intact tissues and dense 3D cultures, PLOS Biology 17 (8) (2019) e3000388. doi:10.1371/journal.pbio.3000388. URL https://dx.plos.org/10.1371/journal.pbio.3000388

4. S. Berg, D. Kutra, T. Kroeger, C. N. Straehle, B. X. Kausler, C. Haubold, M. Schiegg, J. Ales, T. Beier,M. Rudy, K. Eren, J. I. Cervantes, B. Xu, F. Beuttenmueller, A. Wolny, C. Zhang, U. Koethe, F. A. Hamprecht, A. Kreshuk, ilastik: interactive machine learning for (bio)image analysis, Nature Methods 16 (12) (2019) 1226–1232. doi:10.1038/s41592-019-0582-9. URL http://www.nature.com/articles/s41592-019-0582-9

5. J. C. Caicedo, A. Goodman, K. W. Karhohs, B. A. Cimini, J. Ackerman, M. Haghighi, C. Heng, T. Becker, M. Doan, C. McQuin, M. Rohban, S. Singh, A. E. Carpenter, Nucleus segmentation across imaging experiments: the 2018 Data Science Bowl, Nature Methods 16 (12) (2019) 1247–1253. doi:10.1038/s41592-019-0612-7. URL http://www.nature.com/articles/s41592-019-0612-7

6. Y. Tokuoka, T. G. Yamada, D. Mashiko, Z. Ikeda, N. F. Hiroi, T. J. Kobayashi, K. Yamagata Funahashi, 3D convolutional neural networks-based segmentation to acquire quantitative criteria of the nucleus during mouse embryogenesis, npj Systems Biology and Applications 6 (1) (2020) 32. A. doi:10.1038/s41540-020-00152-8. URL https://www.nature.com/articles/s41540-020-00152-8

7. E. Posfai, S. Petropoulos, F. R. O. de Barros, J. P. Schell, I. Jurisica, R. Sandberg, F. Lanner, J. Rossant, Position- and Hippo signaling-dependent plasticity during lineage segregation in the early mouse embryo, eLife 6 (2017) e22906. doi:10.7554/eLife.22906. URL https://elifesciences.org/articles/22906

8. J. Deng, W. Dong, R. Socher, L.-J. Li, Kai Li, Li Fei-Fei, ImageNet: A large-scale hierarchical image database, in: 2009 IEEE Conference on Computer Vision and Pattern Recognition, IEEE, Miami, FL, 2009, pp. 248–255. doi:10.1109/CVPR.2009.5206848. URL https://ieeexplore.ieee.org/document/5206848/

9. G. De Medeiros, R. Ortiz, P. Strnad, A. Boni, F. Moos, N. Repina, L. Challet Meylan, F. Maurer, P. Liberali, Multiscale light-sheet organoid imaging framework, Nature Communications 13 (1) (2022) 4864. doi:10.1038/s41467-022-32465-z. URL https://www.nature.com/articles/s41467-022-32465-z

10. M. Lalit, P. Tomancak, F. Jug, Embedseg: Embedding-based instance segmentation for biomedical microscopy data, Medical Image Analysis 81 (2022) 102523. doi:https://doi.org/10.1016/j.media.2022.102523. URL https://www.sciencedirect.com/science/article/pii/S1361841522001700

11. B. Gu, E. Posfai, J. Rossant, Efficient generation of targeted large insertions by microinjection into two-cell-stage mouse embryos, Nature Biotechnology 36 (7) (2018) 632–637. doi:10.1038/nbt.4166. URL http://www.nature.com/articles/nbt.4166

12. J.-P. Concordet, M. Haeussler, CRISPOR: intuitive guide selection for CRISPR/Cas9 genome editing experiments and screens, Nucleic Acids Research 46 (W1) (2018) W242–W245. arXiv:https://academic.oup.com/nar/article-pdf/46/W1/W242/25110393/gky354.pdf, doi:10.1093/nar/gky354. URL https://doi.org/10.1093/nar/gky354

13. B. Gu, E. Posfai, M. Gertsenstein, J. Rossant, Efficient generation of large-fragment knock-in mouse models using 2-cell (2c)-homologous recombination (hr)-crispr, Current Protocols in Mouse Biology 10 (1) (2020) e67. arXiv:https://currentprotocols.onlinelibrary.wiley.com/doi/pdf/10.1002/cpmo.67, doi:10.1002/cpmo.67. URL https://currentprotocols.onlinelibrary.wiley.com/doi/abs/10.1002/cpmo.67

14. K. McDole, Y. Zheng, Generation and live imaging of an endogenous Cdx2 reporter mouse line, genesis 50 (10) (2012) 775–782. doi:10.1002/dvg.22049. URL https://onlinelibrary.wiley.com/doi/10.1002/dvg.22049

15. M. D. Muzumdar, B. Tasic, K. Miyamichi, L. Li, L. Luo, A global double-fluorescent Cre reporter mouse, Genesis (New York, N.Y.: 2000) 45 (9) (2007) 593–605. doi:10.1002/dvg.20335.

16. B. Gu, B. Bradshaw, M. Zhu, Y. Sun, S. Hopyan, J. Rossant, Live imaging YAP signalling in mouse embryo development, Open Biology 12 (1) (2022) 210335. doi:10.1098/rsob.210335.

17. D. Hirling, E. Tasnadi, J. Caicedo, M. V. Caroprese, R. Sjögren, M. Aubreville, K. Koos, P. Horvath, Segmentation metric misinterpretations in bioimage analysis, Nature Methods (Jul. 2023). doi:10.1038/s41592-023-01942-8. URL https://www.nature.com/articles/s41592-023-01942-8

18. R. Hollandi, A. Diosdi, G. Hollandi, N. Moshkov, P. Horvath, AnnotatorJ: an ImageJ plugin to ease hand annotation of cellular compartments, Molecular Biology of the Cell 31 (20) (2020) 2179–2186. doi:10.1091/mbc.E20-02-0156. URL https://www.molbiolcell.org/doi/10.1091/mbc.E20-02-0156

19. A. Myronenko, X. Song, Point set registration: Coherent point drift, IEEE Transactions on Pattern Analysis and Machine Intelligence 32 (12) (2010) 2262–2275. doi:10.1109/TPAMI.2010.46.

20. P. Strnad, S. Gunther, J. Reichmann, U. Krzic, B. Balazs, G. de Medeiros, N. Norlin, T. Hiiragi, L. Hufnagel, J. Ellenberg, Inverted light-sheet microscope for imaging mouse pre-implantation development, Nature Methods 13 (2) (2016) 139–142. doi:10.1038/nmeth.3690.

21. R. Niwayama, P. Moghe, Y.-J. Liu, D. Fabrèges, F. Buchholz, M. Piel, T. Hiiragi, A Tug-of-War between Cell Shape and Polarity Controls Division Orientation to Ensure Robust Patterning in the Mouse Blastocyst, Developmental Cell 51 (5) (2019) 564–574.e6. doi:10.1016/j.devcel.2019.10.012. URL https://linkinghub.elsevier.com/retrieve/pii/S1534580719308561

22. P. P. Laissue, R. A. Alghamdi, P. Tomancak, E. G. Reynaud, H. Shroff, Assessing phototoxicity in live fluorescence imaging, Nature Methods 14 (7) (2017) 657–661. doi:10.1038/nmeth.4344. URL http://www.nature.com/articles/nmeth.4344

23. C. Stringer, T. Wang, M. Michaelos, M. Pachitariu, Cellpose: a generalist algorithm for cellular segmentation, Nature Methods 18 (1) (2021) 100–106. doi:10.1038/s41592-020-01018-x. URL http://www.nature.com/articles/s41592-020-01018-x

24. M. Weigert, U. Schmidt, R. Haase, K. Sugawara, G. Myers, Star-convex polyhedra for 3d object detection and segmentation in microscopy, in: The IEEE Winter Conference on Applications of Computer Vision (WACV), 2020. doi:10.1109/WACV45572.2020.9093435.

25. R. Ortiz, G. de Medeiros, A. H. F. M. Peters, P. Liberali, M. Rempfler, RDCNet: Instance Segmentation with a Minimalist Recurrent Residual Network, in: M. Liu, P. Yan, C. Lian, X. Cao (Eds.), Machine Learning in Medical Imaging, Vol. 12436, Springer International Publishing, Cham, 2020, pp. 434–443. doi:10.1007/978-3-030-59861-7_44. URL https://link.springer.com/10.1007/978-3-030-59861-7_44

26. Z. Lin, D. Wei, M. D. Petkova, Y. Wu, Z. Ahmed, K. S. K S. Zou, N. Wendt, J. Boulanger-Weill, X. Wang, N. Dhanyasi, I. Arganda-Carreras, F. Engert, J. Lichtman, H. Pfister, NucMM Dataset: 3D Neuronal Nuclei Instance Segmentation at Sub-Cubic Millimeter Scale, in: M. de Bruijne, P. C. Cattin, S. Cotin, N. Padoy, S. Speidel, Y. Zheng, C. Essert (Eds.), Medical Image Computing and Computer Assisted Intervention – MICCAI 2021, Vol. 12901, Springer International Publishing, Cham, 2021, pp. 164–174. doi:10.1007/978-3-030-87193-2_16. URL https://link.springer.com/10.1007/978-3-030-87193-2_16

27. A. Hatamizadeh, Y. Tang, V. Nath, D. Yang, A. Myronenko, B. Landman, H. R. Roth, D. Xu, UNETR: Transformers for 3D Medical Image Segmentation, in: 2022 IEEE/CVF Winter Conference on Applications of Computer Vision (WACV), IEEE, Waikoloa, HI, USA, 2022, pp. 1748–1758. doi:10.1109/WACV51458.2022.00181. URL https://ieeexplore.ieee.org/document/9706678/

28. K. Sugawara, C. Cevrim, M. Averof, Tracking cell lineages in 3D by incremental deep learning, eLife 11 (2022) e69380. doi:10.7554/eLife.69380. URL https://elifesciences.org/articles/69380

29. N. Saiz, K. M. Williams, V. E. Seshan, A.-K. Hadjantonakis, Asynchronous fate decisions by single cells collectively ensure consistent lineage composition in the mouse blastocyst, Nature Communications 7 (1) (2016) 13463. doi:10.1038/ncomms13463. URL http://www.nature.com/articles/ncomms13463

30. S. Syed, H. Wilky, J. Raimundo, B. Lim, A. A. Amodeo, The nuclear to cytoplasmic ratio directly regulates zygotic transcription in Drosophila through multiple modalities, Proceedings of the National Academy of Sciences 118 (14) (2021) e2010210118. doi:10.1073/pnas.2010210118. URL https://pnas.org/doi/full/10.1073/pnas.2010210118

31. S. Balachandra, S. Sarkar, A. A. Amodeo, The Nuclear-to-Cytoplasmic Ratio: Coupling DNA Content to Cell Size, Cell Cycle, and Biosynthetic Capacity, Annual Review of Genetics 56 (2022) 165–185. doi:10.1146/annurev-genet-080320-030537.

32. C. E. M. Aiken, P. P. L. Swoboda, J. N. Skepper, M. H. Johnson, The direct measurement of embryogenic volume and nucleo-cytoplasmic ratio during mouse pre-implantation development, Reproduction 128 (5) (2004) 527–535. doi:10.1530/rep.1.00281. URL https://rep.bioscientifica.com/view/journals/rep/128/5/1280527.xml

33. E. Tsichlaki, G. FitzHarris, Nucleus downscaling in mouse embryos is regulated by cooperative developmental and geometric programs, Scientific Reports 6 (1) (2016) 28040. doi:10.1038/srep28040. URL http://www.nature.com/articles/srep28040

34. C. J. Chan, M. Costanzo, T. Ruiz-Herrero, G. Mönke, R. J. Petrie, M. Bergert, A. Diz-Muñoz, L. Mahadevan, T. Hiiragi, Hydraulic control of mammalian embryo size and cell fate, Nature 571 (7763) (2019) 112–116. doi:10.1038/s41586-019-1309-x.

35. P. Moghe, R. Belousov, T. Ichikawa, C. Iwatani, T. Tsukiyama, F. Graner, A. Erzberger, T. Hiiragi, Apical-driven cell sorting optimised for tissue geometry ensures robust patterning, bioRxiv (2023). arXiv:https://www.biorxiv.org/content/early/2023/05/17/2023.05.16.540918.full.pdf, doi:10.1101/2023.05.16.540918. URL https://www.biorxiv.org/content/early/2023/05/17/2023.05.16.540918

36. A. Hallou, H. G. Yevick, B. Dumitrascu, V. Uhlmann, Deep learning for bioimage analysis in developmental biology, Development 148 (18) (2021) dev199616. doi:10.1242/dev.199616. URL https://journals.biologists.com/dev/article/148/18/dev199616/272084/Deep-learning-for-bioimage-analysis-in

37. N. Koushki, A. Ghagre, L. K. Srivastava, C. Sitaras, H. Yoshie, C. Molter, A. J. Ehrlicher, Lamin a redistribution mediated by nuclear deformation determines dynamic localization of yap, bioRxiv (2020). arXiv:https://www.biorxiv.org/content/early/2020/03/20/2020.03.19.998708.full.pdf, doi:10.1101/2020.03.19.998708. URL https://www.biorxiv.org/content/early/2020/03/20/2020.03.19.998708

38. W. Ouyang, F. Beuttenmueller, E. Gómez-de Mariscal, C. Pape, T. Burke, C. Garcia-López-de Haro, C. Russell, L. Moya-Sans, C. de-la Torre-Gutiérrez, D. Schmidt, D. Kutra, M. Novikov, M. Weigert, U. Schmidt, P. Bankhead, G. Jacquemet, D. Sage, R. Henriques, A. Muñoz-Barrutia, E. Lundberg, F. Jug, A. Kreshuk, Bioimage model zoo: A community-driven resource for accessible deep learning in bioimage analysis, bioRxiv (2022). arXiv:https://www.biorxiv.org/content/early/2022/06/08/2022.06.07.495102.full.pdf, doi:10.1101/2022.06.07.495102. URL https://www.biorxiv.org/content/early/2022/06/08/2022.06.07.495102

39. L. Yang, R. P. Ghosh, J. M. Franklin, S. Chen, C. You, R. R. Narayan, M. L. Melcher, J. T. Liphardt, NuSeT: A deep learning tool for reliably separating and analyzing crowded cells, PLOS Computational Biology 16 (9) (2020) e1008193. doi:10.1371/journal.pcbi.1008193. URL https://dx.plos.org/10.1371/journal.pcbi.1008193

40. O. Cicek, A. Abdulkadir, S. S. Lienkamp, T. Brox, O. Ronneberger, 3D U-Net: Learning Dense Volumetric Segmentation from Sparse Annotation, in: S. Ourselin, L. Joskowicz, M. R. Sabuncu, G. Unal, W. Wells (Eds.), Medical Image Computing and Computer-Assisted Intervention – MICCAI 2016, Vol. 9901, Springer International Publishing, Cham, 2016, pp. 424–432. doi:10.1007/978-3-319-46723-8_49. URL https://link.springer.com/10.1007/978-3-319-46723-8_49

41. V. Ulman, M. Maska, K. E. G. Magnusson, O. Ronneberger, C. Haubold, N. Harder, P. Matula, P. Matula, D. Svoboda, M. Radojevic, I. Smal, K. Rohr, J. Jalden, H. M. Blau, O. Dzyubachyk, B. Lelieveldt, P. Xiao, Y. Li, S.-Y. Cho, A. C. Dufour, J.-C. Olivo-Marin, C. C. Reyes-Aldasoro, J. A. Solis-Lemus, R. Bensch, T. Brox, J. Stegmaier, R. Mikut, S. Wolf, F. A. Hamprecht, T. Esteves, P. Quelhas, O. Demirel, L. Malmström, F. Jug, P. Tomancak, E. Meijering, A. Munoz-Barrutia, M. Kozubek, C. Ortiz-de Solorzano, An objective comparison of cell-tracking algorithms, Nature Methods 14 (12) (2017) 1141–1152. doi:10.1038/nmeth.4473. URL http://www.nature.com/articles/nmeth.4473

42. V. Bondarenko, M. Nikolaev, D. Kromm, R. Belousov, A. Wolny, S. Rezakhani, J. Hugger, V. Uhlmann, L. Hufnagel, A. Kreshuk, J. Ellenberg, A. Erzberger, M. Lutolf, T. Hiiragi, Coordination between embryo growth and trophoblast migration upon implantation delineates mouse embryogenesis, bioRxiv (2022). arXiv:https://www.biorxiv.org/content/early/2022/06/16/2022.06.13.495767.full.pdf, doi:10.1101/2022.06.13.495767. URL https://www.biorxiv.org/content/early/2022/06/16/2022.06.13.495767

43. T.-Y. Lin, M. Maire, S. Belongie, J. Hays, P. Perona, D. Ramanan, P. Dollár, C. L. Zitnick, Microsoft COCO: Common Objects in Context, in: D. Fleet, T. Pajdla, B. Schiele, T. Tuytelaars (Eds.), Computer Vision – ECCV 2014, Vol. 8693, Springer International Publishing, Cham, 2014, pp. 740–755. doi:10.1007/978-3-319-10602-1_48. URL http://link.springer.com/10.1007/978-3-319-10602-1_48

