## Supplemental figures for "Nuclear instance segmentation and tracking for preimplantation mouse embryos"

Fig S1

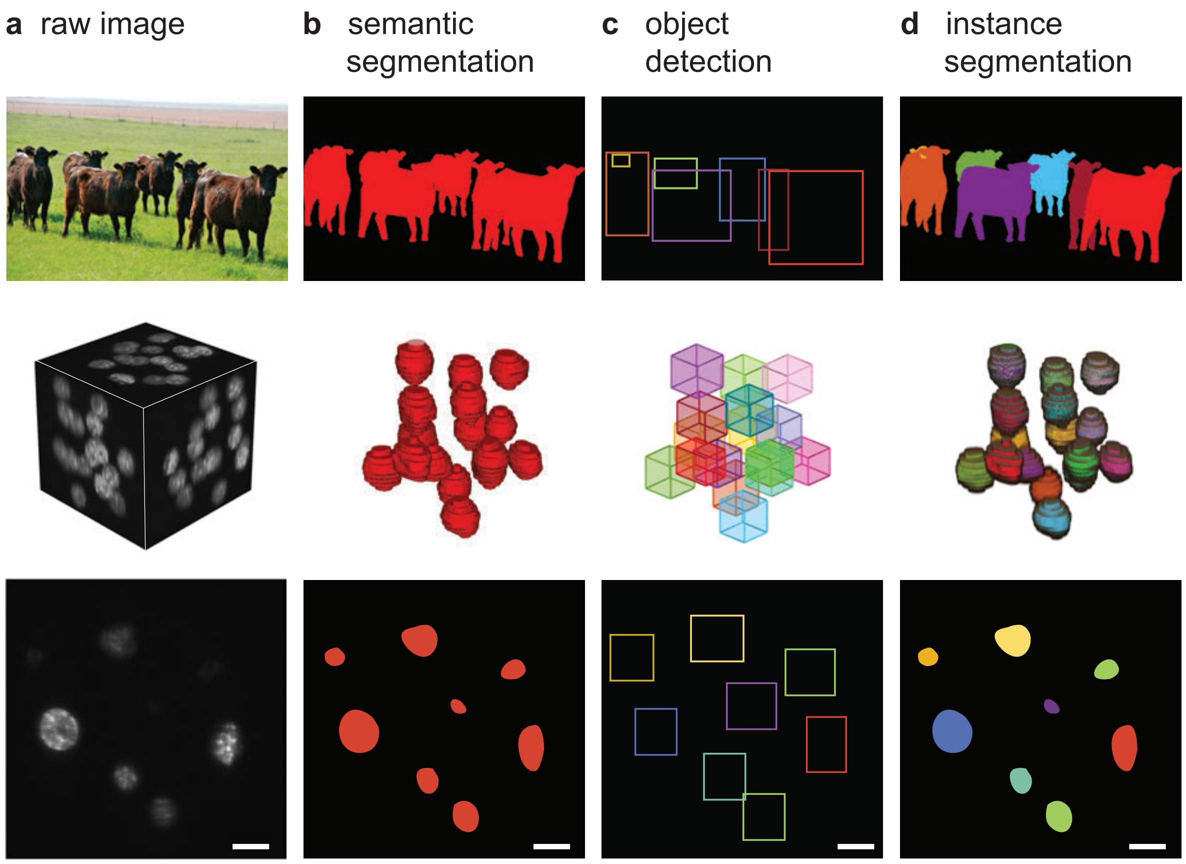
a raw image
b semantic segmentation
c object detection
d instance segmentation

Fig S2

A

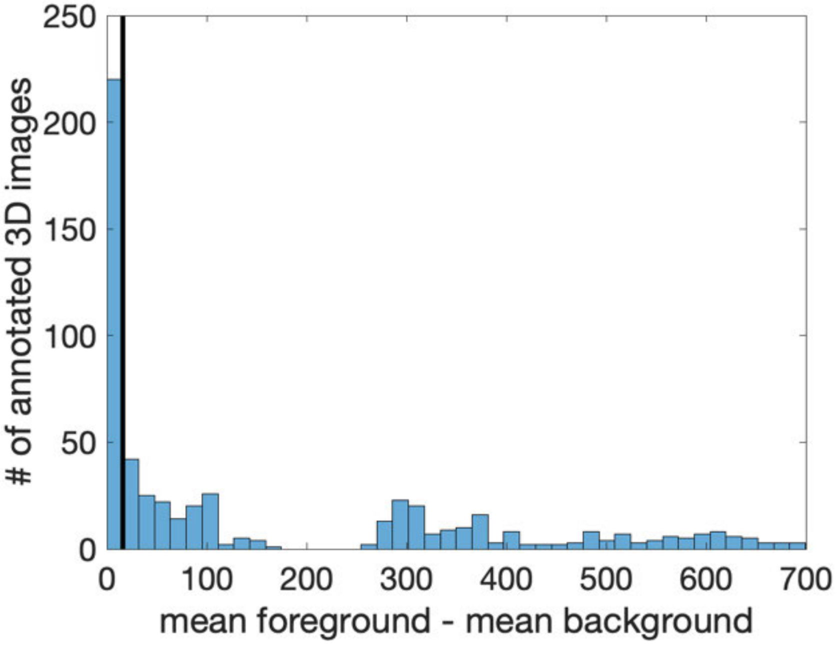

250
200
150
100
50
0
0
100
200
300
400
500
600
700
### of annotated 3D images
mean foreground - mean background


B

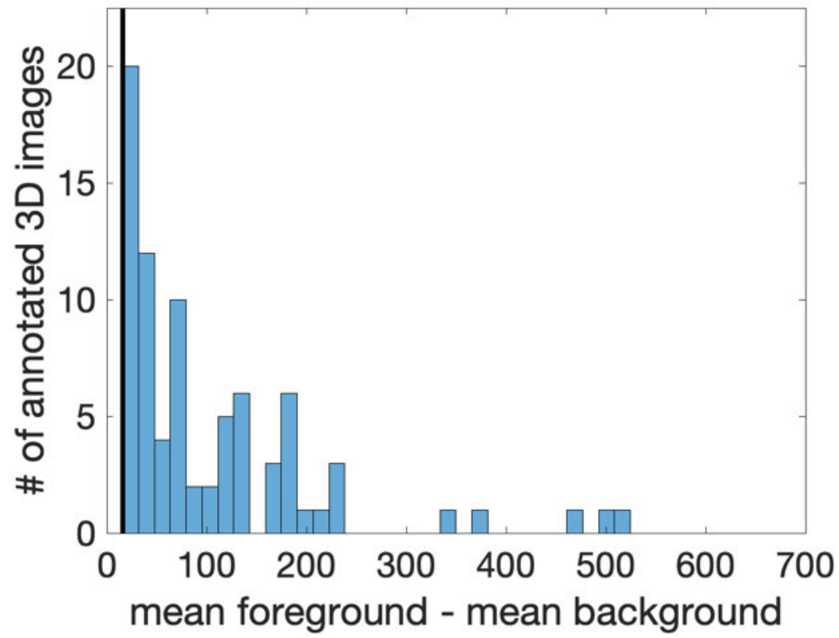

20
15
10
5
0
0
100
200
300
400
500
600
700
### of annotated 3D images
mean foreground - mean background

Fig S3

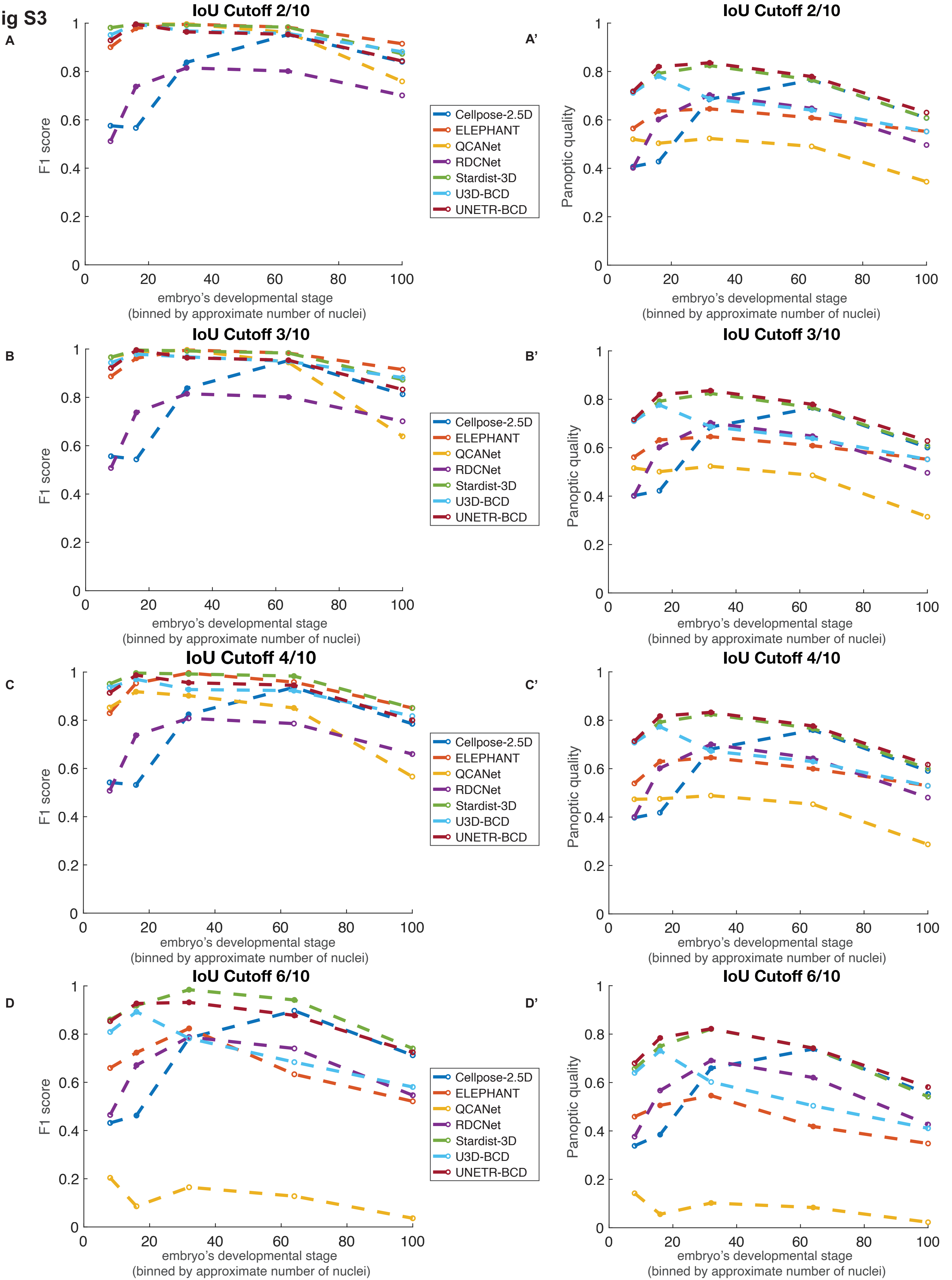

A
IoU Cutoff 2/10
F1 score
embryo's developmental stage
(binned by approximate number of nuclei)
Cellpose-2.5D
ELEPHANT
QCANet
RDCNet
Stardist-3D
U3D-BCD
UNETR-BCD
A'
IoU Cutoff 2/10
Panoptic quality
embryo's developmental stage
(binned by approximate number of nuclei)
B
IoU Cutoff 3/10
F1 score
embryo's developmental stage
(binned by approximate number of nuclei)
Cellpose-2.5D
ELEPHANT
QCANet
RDCNet
Stardist-3D
U3D-BCD
UNETR-BCD
B'
IoU Cutoff 3/10
Panoptic quality
embryo's developmental stage
(binned by approximate number of nuclei)
C
IoU Cutoff 4/10
F1 score
embryo's developmental stage
(binned by approximate number of nuclei)
Cellpose-2.5D
ELEPHANT
QCANet
RDCNet
Stardist-3D
U3D-BCD
UNETR-BCD
C'
IoU Cutoff 4/10
Panoptic quality
embryo's developmental stage
(binned by approximate number of nuclei)
D
IoU Cutoff 6/10
F1 score
embryo's developmental stage
(binned by approximate number of nuclei)
Cellpose-2.5D
ELEPHANT
QCANet
RDCNet
Stardist-3D
U3D-BCD
UNETR-BCD
D'
IoU Cutoff 6/10
Panoptic quality
embryo's developmental stage
(binned by approximate number of nuclei)

Fig S4

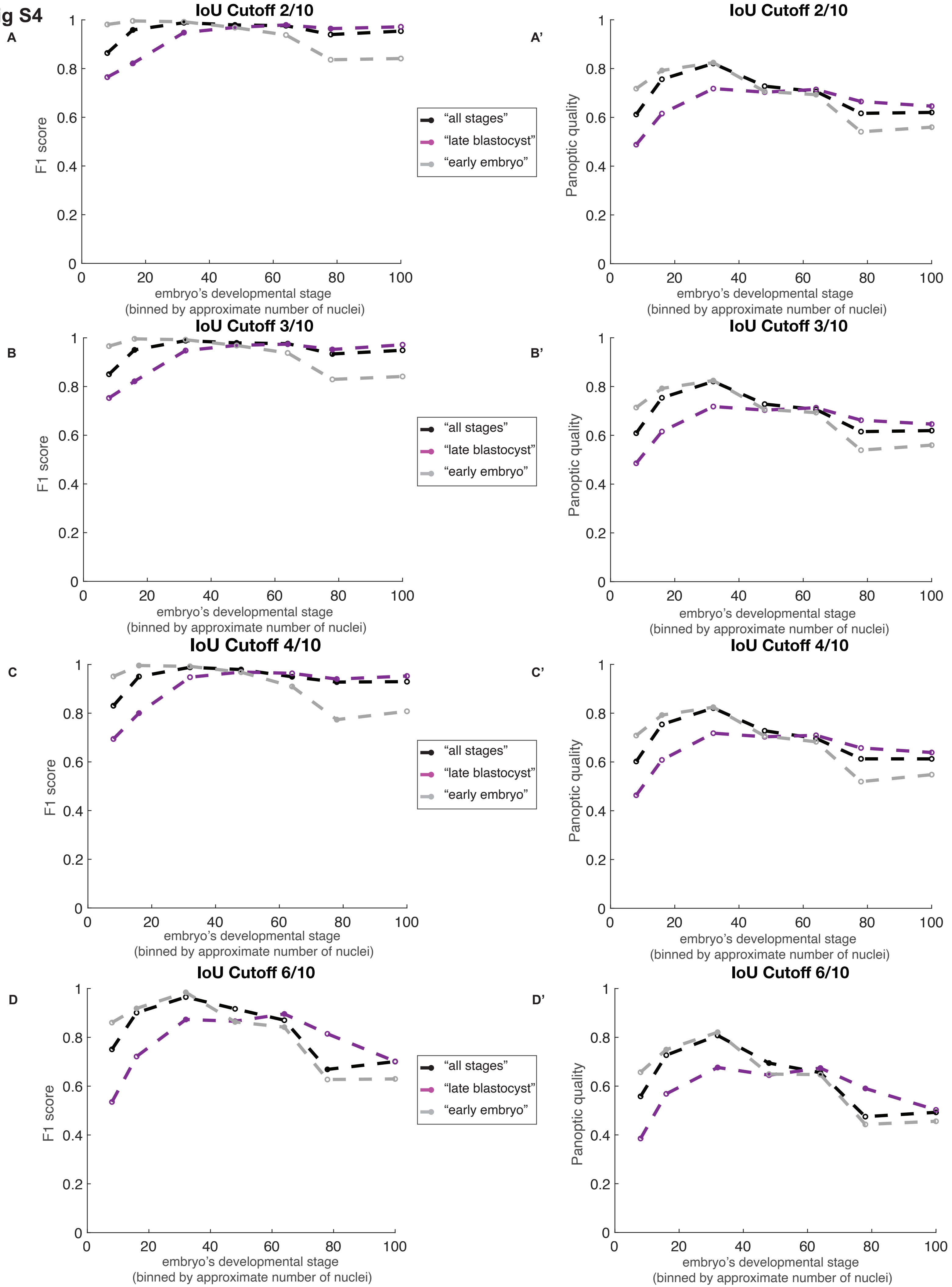

A
IoU Cutoff 2/10
F1 score
embryo's developmental stage
(binned by approximate number of nuclei)
"all stages"
"late blastocyst"
"early embryo"
A'
IoU Cutoff 2/10
Panoptic quality
embryo's developmental stage
(binned by approximate number of nuclei)
B
IoU Cutoff 3/10
F1 score
embryo's developmental stage
(binned by approximate number of nuclei)
"all stages"
"late blastocyst"
"early embryo"
B'
IoU Cutoff 3/10
Panoptic quality
embryo's developmental stage
(binned by approximate number of nuclei)
C
IoU Cutoff 4/10
F1 score
embryo's developmental stage
(binned by approximate number of nuclei)
"all stages"
"late blastocyst"
"early embryo"
C'
IoU Cutoff 4/10
Panoptic quality
embryo's developmental stage
(binned by approximate number of nuclei)
D
IoU Cutoff 6/10
F1 score
embryo's developmental stage
(binned by approximate number of nuclei)
"all stages"
"late blastocyst"
"early embryo"
D'
IoU Cutoff 6/10
Panoptic quality
embryo's developmental stage
(binned by approximate number of nuclei)

Fig S5

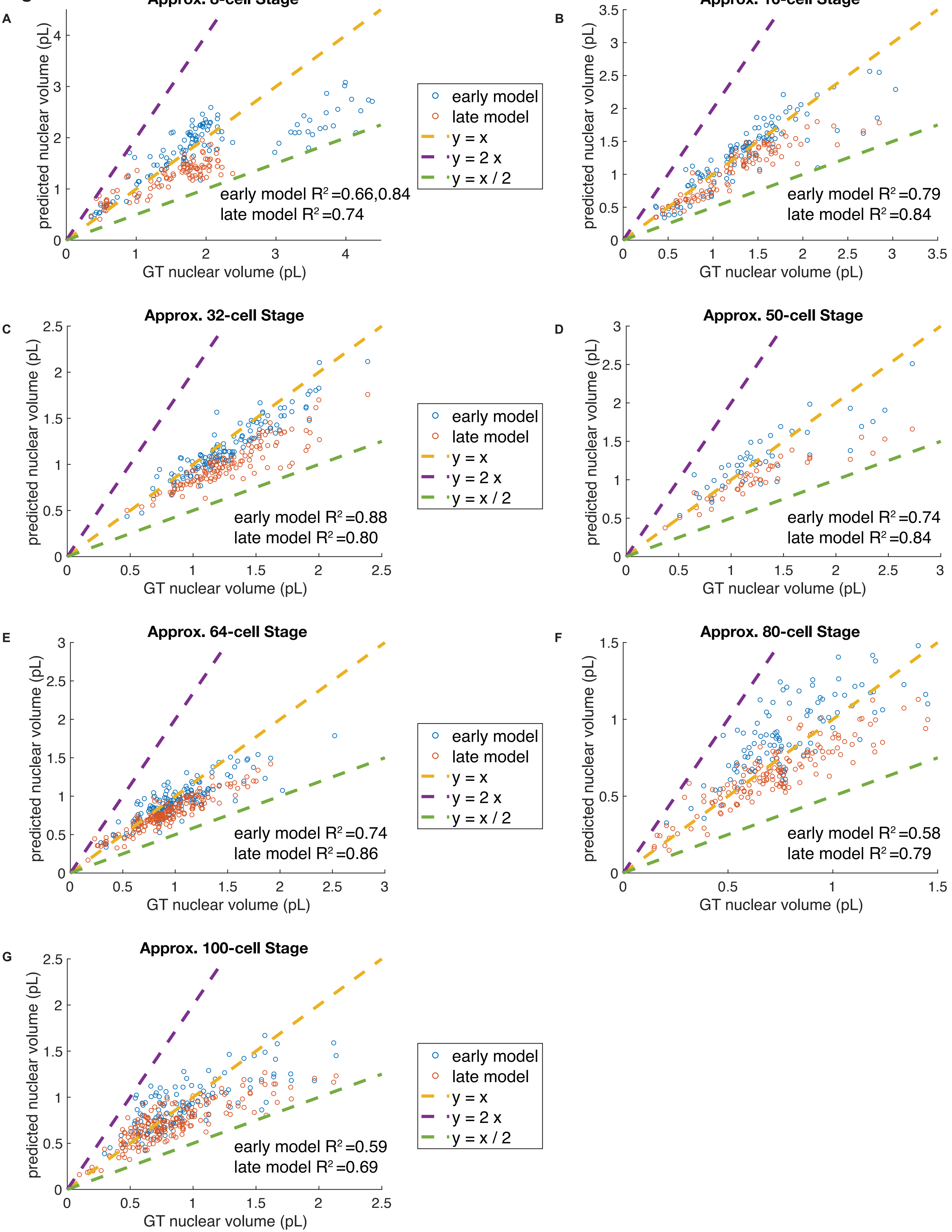
A
Approx. 8-cell Stage
predicted nuclear volume (pL)
GT nuclear volume (pL)
early model
late model
y = x
y = 2 x
y = x / 2
early model R² =0.66,0.84
late model R² =0.74
B
Approx. 16-cell Stage
early model R² =0.79
late model R² =0.84
C
Approx. 32-cell Stage
early model R² =0.88
late model R² =0.80
D
Approx. 50-cell Stage
early model R² =0.74
late model R² =0.84
E
Approx. 64-cell Stage
early model R² =0.74
late model R² =0.86
F
Approx. 80-cell Stage
early model R² =0.58
late model R² =0.79
G
Approx. 100-cell Stage
early model R² =0.59
late model R² =0.69

Fig S6

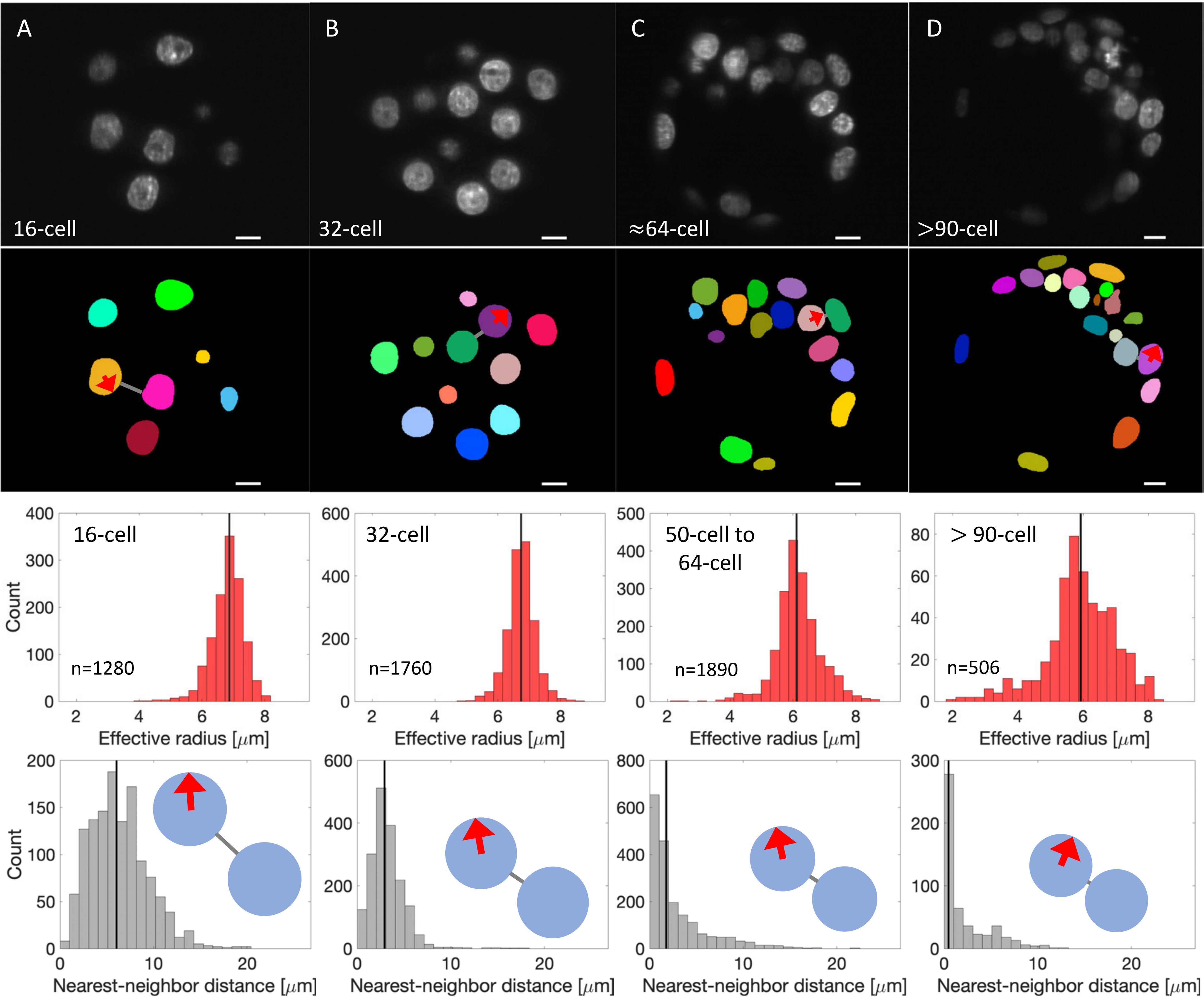

A
B
C
D
16-cell
32-cell
≈64-cell
>90-cell
16-cell
n=1280
32-cell
n=1760
50-cell to
64-cell
n=1890
> 90-cell
n=506
Count
Effective radius [μm]
Count
Nearest-neighbor distance [μm]

Fig S7

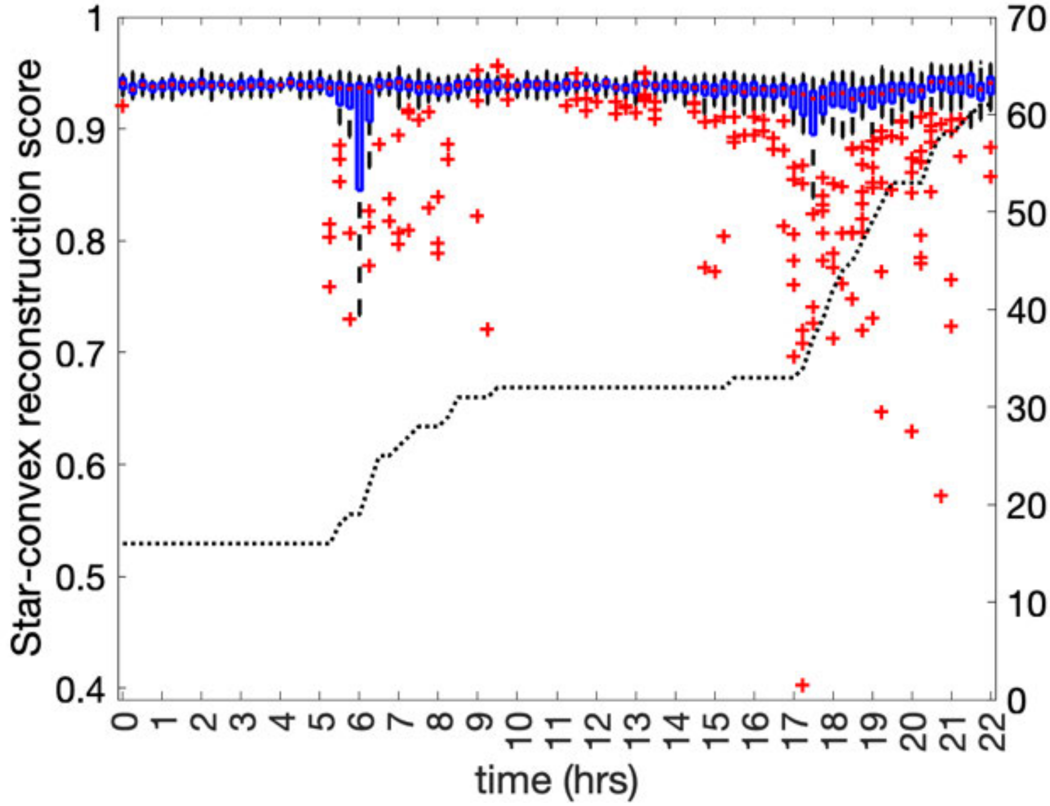

Star-convex reconstruction score
number of nuclei
time (hrs)

**Fig S8**

**A** False positives near bright polar body

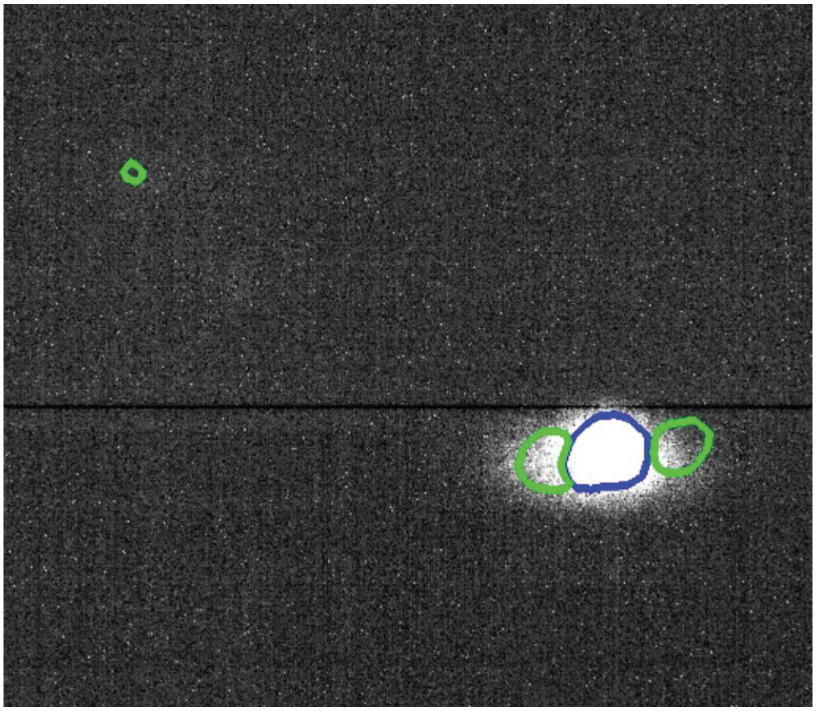

True positive: blue
False positive: green

**B** False negatives induced by debris and reflection from chamber

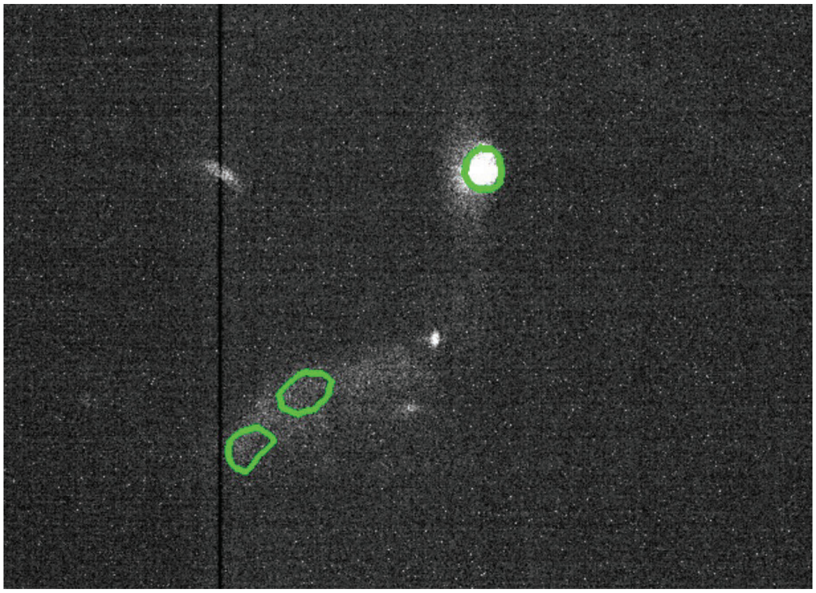

True positive: blue
False positive: green

**C** False negative induced by debris

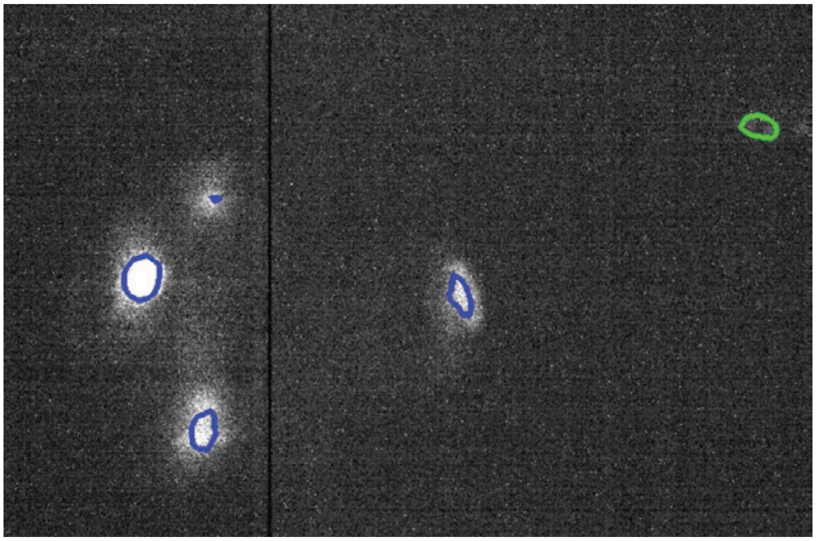

True positive: blue
False positive: green

Fig S9

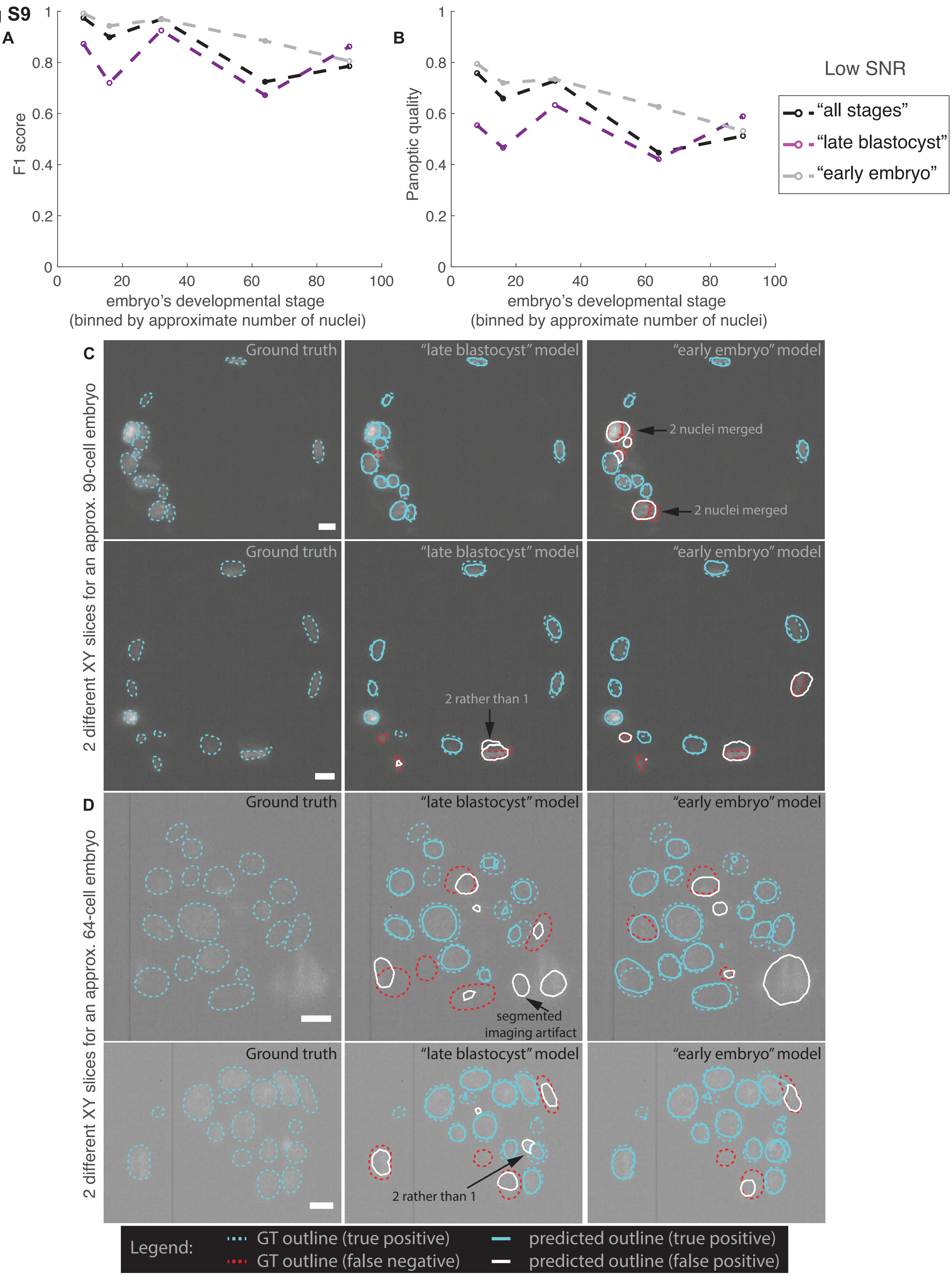
A
F1 score
embryo's developmental stage
(binned by approximate number of nuclei)
B
Panoptic quality
embryo's developmental stage
(binned by approximate number of nuclei)
Low SNR
"all stages"
"late blastocyst"
"early embryo"
C
2 different XY slices for an approx. 90-cell embryo
Ground truth
"late blastocyst" model
"early embryo" model
2 nuclei merged
2 nuclei merged
2 rather than 1
D
2 different XY slices for an approx. 64-cell embryo
Ground truth
"late blastocyst" model
"early embryo" model
segmented imaging artifact
2 rather than 1
Legend:
GT outline (true positive)
GT outline (false negative)
predicted outline (true positive)
predicted outline (false positive)

Fig S10

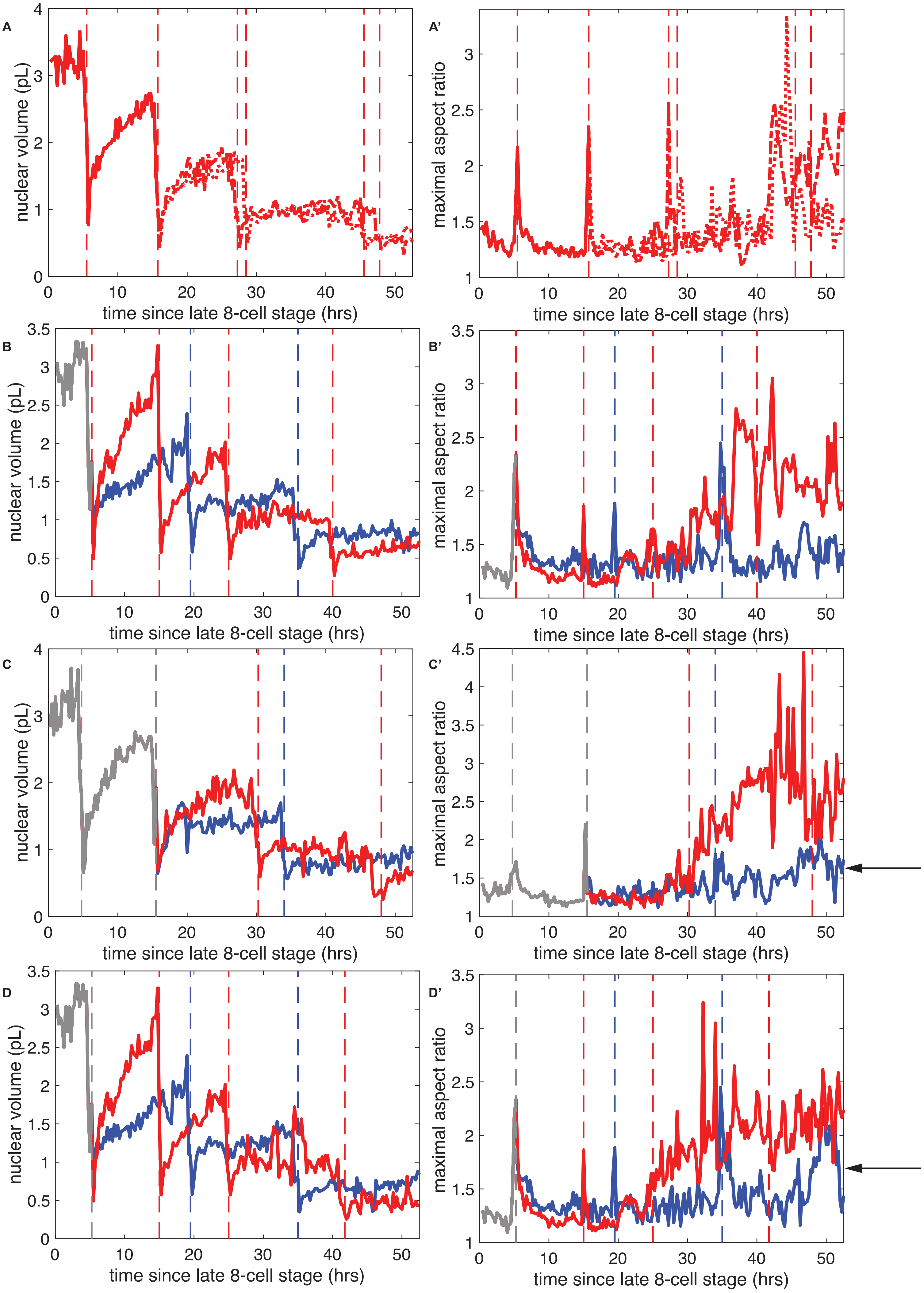

A
A'
B
B'
C
C'
D
D'
nuclear volume (pL)
maximal aspect ratio
time since late 8-cell stage (hrs)
0
10
20
30
40
50

Fig S11

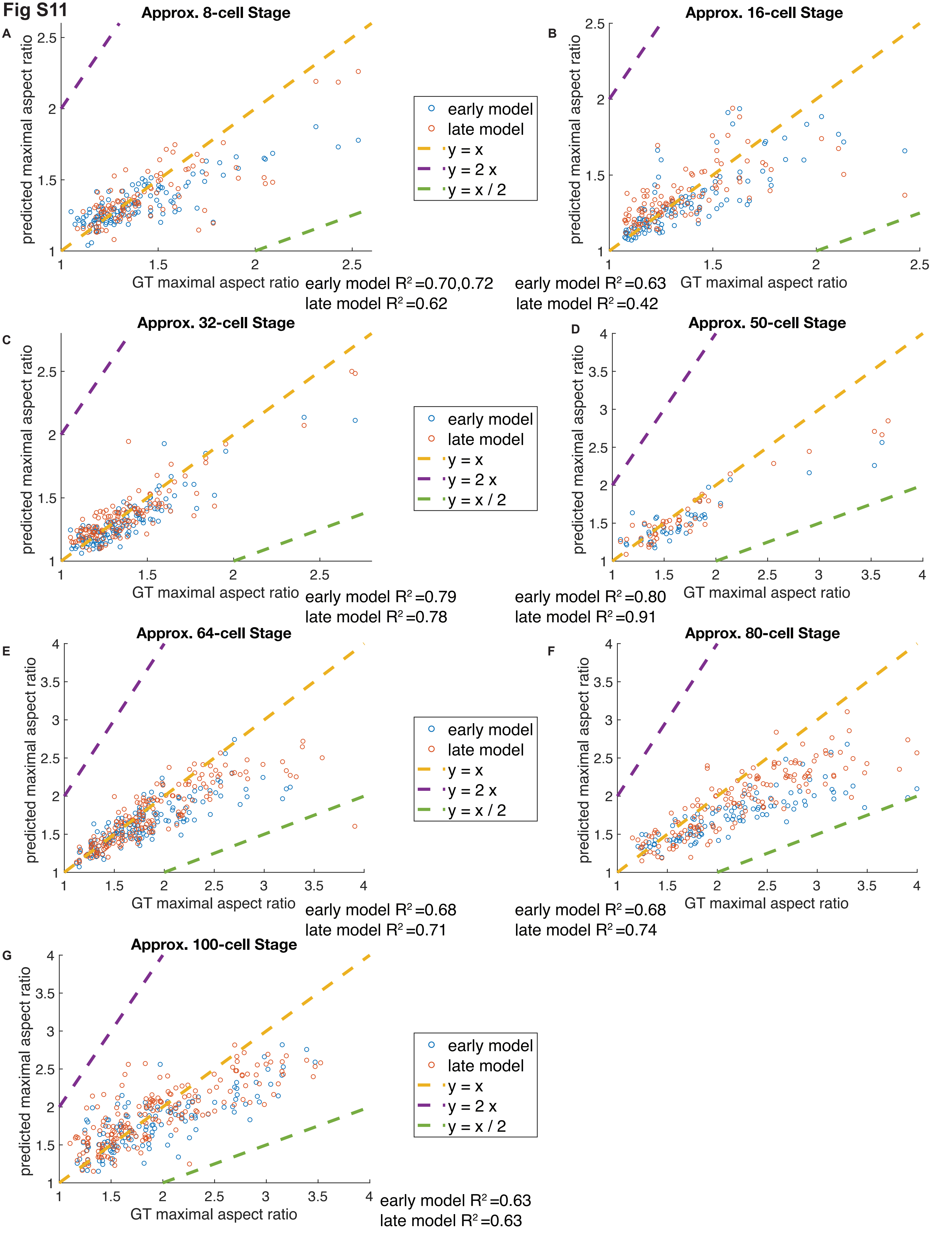

A
Approx. 8-cell Stage
early model R$^2$=0.70,0.72
late model R$^2$=0.62
B
Approx. 16-cell Stage
early model R$^2$=0.63
late model R$^2$=0.42
C
Approx. 32-cell Stage
early model R$^2$=0.79
late model R$^2$=0.78
D
Approx. 50-cell Stage
early model R$^2$=0.80
late model R$^2$=0.91
E
Approx. 64-cell Stage
early model R$^2$=0.68
late model R$^2$=0.71
F
Approx. 80-cell Stage
early model R$^2$=0.68
late model R$^2$=0.74
G
Approx. 100-cell Stage
early model R$^2$=0.63
late model R$^2$=0.63
predicted maximal aspect ratio
GT maximal aspect ratio
early model
late model
y = x
y = 2 x
y = x / 2

#### Fig S12

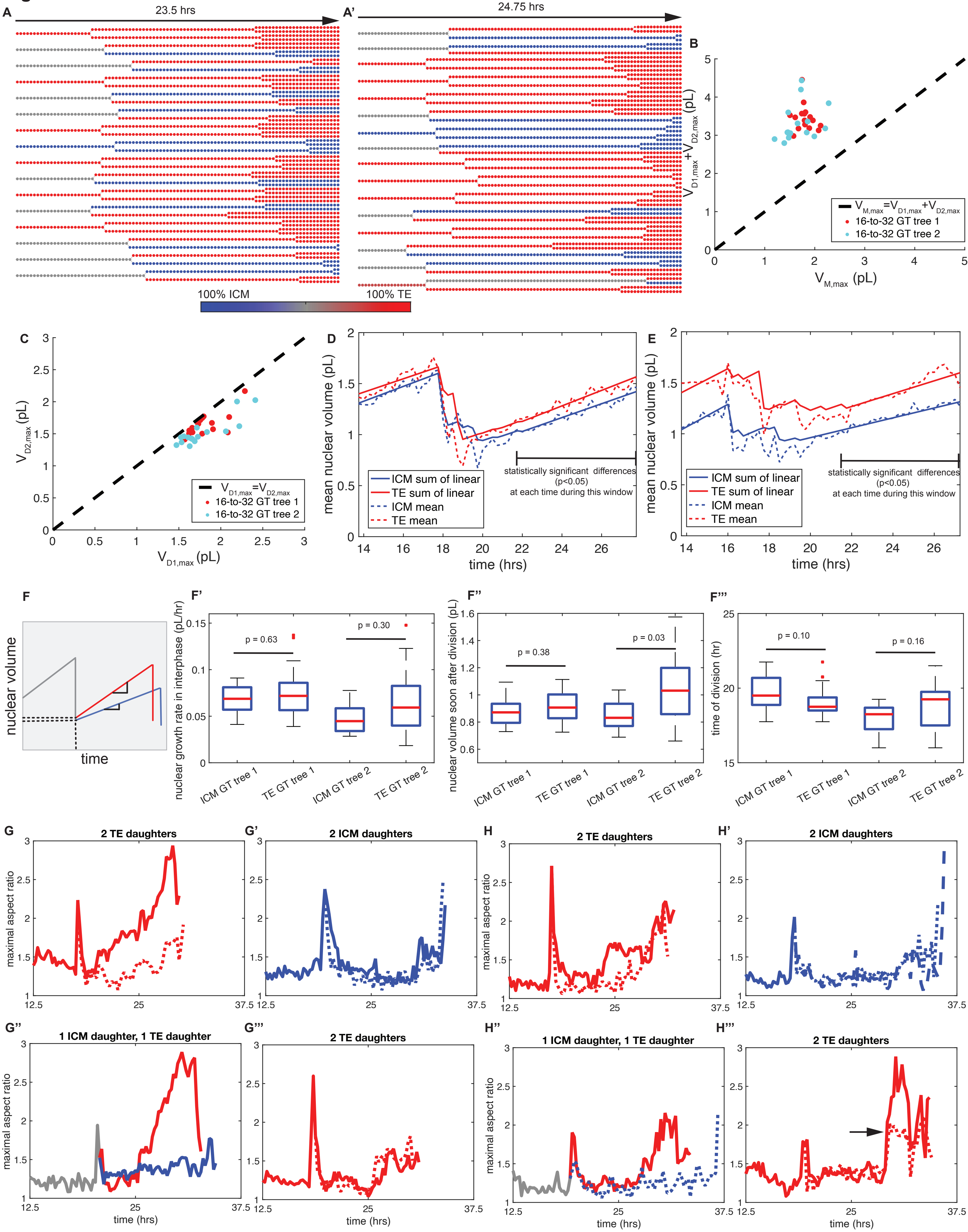

A
23.5 hrs
A'
24.75 hrs
100% ICM
100% TE
B
$V_{D1,max}+V_{D2,max}$ (pL)
$V_{M,max}$ (pL)
$V_{M,max}=V_{D1,max}+V_{D2,max}$
16-to-32 GT tree 1
16-to-32 GT tree 2
C
$V_{D2,max}$ (pL)
$V_{D1,max}$ (pL)
$V_{D1,max}=V_{D2,max}$
16-to-32 GT tree 1
16-to-32 GT tree 2
D
mean nuclear volume (pL)
time (hrs)
ICM sum of linear
TE sum of linear
ICM mean
TE mean
statistically significant differences (p<0.05) at each time during this window
E
mean nuclear volume (pL)
time (hrs)
ICM sum of linear
TE sum of linear
ICM mean
TE mean
statistically significant differences (p<0.05) at each time during this window
F
nuclear volume
time
F'
nuclear growth rate in interphase (pL/hr)
p = 0.63
p = 0.30
ICM GT tree 1
TE GT tree 1
ICM GT tree 2
TE GT tree 2
F''
nuclear volume soon after division (pL)
p = 0.38
p = 0.03
ICM GT tree 1
TE GT tree 1
ICM GT tree 2
TE GT tree 2
F'''
time of division (hr)
p = 0.10
p = 0.16
ICM GT tree 1
TE GT tree 1
ICM GT tree 2
TE GT tree 2
G
2 TE daughters
maximal aspect ratio
G'
2 ICM daughters
H
2 TE daughters
maximal aspect ratio
H'
2 ICM daughters
G''
1 ICM daughter, 1 TE daughter
maximal aspect ratio
time (hrs)
G'''
2 TE daughters
time (hrs)
H''
1 ICM daughter, 1 TE daughter
maximal aspect ratio
time (hrs)
H'''
2 TE daughters
time (hrs)
